# Pro-regenerative Extracellular Matrix Hydrogel Prevents and Mitigates Pathological Alterations of Pelvic Muscles Following Birth Injury

**DOI:** 10.1101/2021.05.28.446170

**Authors:** Pamela Duran, Francesca Boscolo Sesillo, Lindsey Burnett, Shawn A. Menefee, Mark Cook, Gisselle Zazueta-Damian, Monika Dzieciatkowska, Emmy Do, Saya French, Manali M. Shah, Clyde Sanvictores, Kirk C. Hansen, Matthew Shtrahman, Karen L. Christman, Marianna Alperin

**Author notes:** **Co-Corresponding Authors**: Karen L. Christman, PhD, Department of Bioengineering, Sanford Consortium for Regenerative Medicine, University of California, San Diego, 2880 Torrey Pines Scenic Drive, La Jolla, CA 92037, Marianna Alperin, MD, MS, University of California, San Diego, Department of Obstetrics, Gynecology, and Reproductive Sciences, 9500 Gilman Drive, La Jolla, CA 92093-0863.

## Abstract

Pelvic floor disorders, which include pelvic organ prolapse, and urinary and fecal incontinence, affect millions of women globally and represent a major public health concern. Pelvic floor muscle (PFM) dysfunction has been identified as one of the leading risk factors for the development of these morbid conditions. Even though childbirth, specifically vaginal delivery, has been long recognized as the most important potentially modifiable risk factor for PFM injury, the precise mechanisms of PFM dysfunction following childbirth remain elusive. In this study we demonstrate that PFMs undergo atrophy and severe fibrosis in parous women with symptomatic pelvic organ prolapse compared to age-matched nulliparous cadaveric donors without history of pelvic floor disorders. These pathological alterations are recapitulated in the pre-clinical rat model of simulated birth injury. The transcriptional signature of PFMs post-injury demonstrates a sustained inflammatory response, impairment in muscle anabolism, and persistent expression of extracellular matrix (ECM) remodeling genes. Next, we evaluated the administration of acellular injectable skeletal muscle extracellular matrix hydrogel for the prevention and mitigation of these pathological alterations. Treatment of PFMs with the biomaterial either at the time of birth injury or 4 weeks post-injury reduced muscle atrophy and mitigated fibrotic degeneration. By evaluating gene expression, we demonstrate that these changes are mainly driven by the hydrogel-induced modulation of the immune response and intramuscular fibrosis, as well as enhancement of the endogenous myogenesis. This work furthers our understanding of PFM birth injury and demonstrates proof-of-concept for a new pragmatic pro-regenerative biomaterial approach for treating injured PFMs.

## Introduction

The prevalence of pelvic floor disorders that disproportionately affect women, including pelvic organ prolapse, and urinary and fecal incontinence, ranges from ~25% in the U.S. and China to ~ 50% of the female population of Australia and Japan (*1–4*). Despite not being life-threating, pelvic floor disorders represent a major public health concern due to their high occurrence, associated morbidities, and economic burden. The pelvic floor structural components support the abdominal and pelvic organs by opposing gravitational forces and intra-abdominal pressure, aid in urinary and fecal continence, and contribute to sexual function and parturition. The complex pelvic supportive system includes the vagina, connective tissues, superficial perineal muscles, and pelvic floor muscles (PFMs) that are comprised of the levator ani complex and the coccygeus muscle. PFM dysfunction has been consistently identified as one of the leading risk factors for the development of symptomatic pelvic floor disorders (*5, 6*).

Multiple epidemiological studies have unequivocally identified vaginal childbirth as the leading risk factor for the PFM injury (*6*). Despite this strong association, the precise mechanisms of PFM dysfunction following vaginal delivery remain elusive (*6*). In addition, alterations in the intrinsic PFM components have not been well characterized in women with pelvic floor disorders. Thus, we first sought to determine the morphological properties of PFMs procured from vaginally parous women with symptomatic pelvic organ prolapse and compare these to control specimens procured from nulliparous cadaveric donors without history of pelvic floor disorders.

During vaginal delivery, PFMs are subjected to mechanical strains that significantly exceed the upper physiological limit (~60%) beyond which appendicular skeletal muscle injury ensues (*7*). Based on computational models of human parturition, the enthesial region of the pubovisceralis portion of the levator ani complex experiences the highest strains, up to 300%, during fetal delivery (*8, 9*). These excessive strains have been presumed to cause radiologically detected avulsions; however, this phenotype is not present in all women with pelvic floor dysfunction or decreased PFM strength (*10, 11*). Due to ethical and technical constraints associated with directly probing PFMs in postpartum women, animal models are essential to study other potential phenotypes of PFM birth injury that are not evident on the conventionally used imaging modalities. We previously validated the rat model for the study of the human PFMs (*12*) and observed that the rat pubocaudalis muscle—analogous to the human pubovisceralis—experiences stretch ratios similar to humans during vaginal distention that simulates fetal crowning and delivery (*13*). These strains led to sarcomere hyperelongation and associated myofibrillar disruption of the rat PFMs (*13*), major causes of mechanical skeletal muscle injury (*14*). Unlike studies of the limb muscles, which identify that an inflammatory cascade following acute injury negatively impacts muscle recovery (*14, 15*), the downstream events subsequent to PFM birth injury have not been determined to date. Thus, in the current study we examined early and delayed PFM responses to simulated birth injury using the rat model. We determined that the morphological alterations observed in the injured rat PFMs applicably modeled pathological changes observed in parous women with symptomatic pelvic floor disorders.

Beyond the lack of fundamental knowledge regarding tissue- and cell-level alterations that govern PFM dysfunction, there are currently minimal strategies to preempt the maladaptive recovery of PFMs following birth injury. PFM exercises have been recommended to compensate for PFM weakness in pregnant and postpartum women. However, this therapy is associated with poor adherence (*16*). Furthermore, the existing treatments for PFM dysfunction and the associated pelvic floor disorders are plagued with high failure rates (*17, 18*), while the only preventative strategy for birth injury - Cesarean section – increases the risk of maternal morbidity and mortality compared to vaginal delivery (*19*). Given the dramatic prevalence of pelvic floor disorders, novel preventative and therapeutic approaches are urgently needed to address this public health problem. One potential tactic is to potentiate constructive remodeling of PFMs following birth injury with the ultimate goal of preventing PFM dysfunction and the resultant pelvic floor disorders. We have previously demonstrated that a decellularized porcine skeletal muscle extracellular matrix (ECM) hydrogel (SKM) promoted constructive muscle remodeling and prevented atrophy in ischemic skeletal muscle through upregulation of pathways related to myogenesis and blood vessel development (*20*). Therefore, our final objective was to investigate the efficacy of SKM, administered at two clinically relevant time points, for the prevention and treatment of the pathological PFM alterations consequent to birth injury.

In this study, we observed substantial PFM atrophy and fibrosis in parous women with pelvic organ prolapse compared to PFMs procured from age-comparable nulliparous cadaveric donors without history of pelvic floor disorders. These pathological alterations, replicated in the simulated birth injury pre-clinical model, appear to be governed by the sustained inflammatory response that negatively effects muscle anabolism and potentiates fibrosis. Finally, the acellular pro-regenerative biomaterial prevented and mitigated the pathological alterations of PFMs by modulating the immune response and enhancing myogenesis, signifying its potential as a new preventative therapy for PFM pathological transformations following birth injury.

## Results

### Pelvic floor muscles in women with symptomatic pelvic organ prolapse demonstrate substantial muscle atrophy and fibrosis

We collected biopsies of the pubovisceralis portion of levator ani from nulliparous cadaveric controls without history of pelvic floor disorders and from parous women, undergoing surgeries for symptomatic pelvic organ prolapse (POP). With respect to race/ethnicity, all cadaveric donors were White/Non-Hispanic; in the POP group - 73.9% of subjects were White/Non-Hispanic, 17.4% were White/Hispanic, and 8.7% were Asian. The two groups did not differ with respect to age (control: 72.0 ± 7.5 y; POP: 69.9 ± 3.1 y, P=0.6) or body mass index (control: 25.4 ± 2.4 kg/m^2^; POP: 27.1 ± 1.1 kg/m^2^, P=0.5). The median parity in the POP group was 3 (1-4). To compare the morphological properties of pubovisceralis, we assessed fiber shape and packing, collagen content (fibrosis), fiber area (atrophy), centralization of nuclei (degeneration-regeneration), and intramuscular fat content (fatty degeneration).

In the POP group, 20% of biopsies contained no myofibers on histological examination. In the remaining biopsies, myofibers exhibited disrupted fiber packing, whereas in the control group - moderate fiber packing density was observed (Fig. 1A). A significant decrease in fiber cross-sectional area was found in the POP group relative to the controls (P<0.0001, Fig. 1B). The smaller fiber size was accompanied by the dramatically increased collagen content in the POP group compared to the controls (P=0.001, Fig. 1C). The proportion of centralized nuclei (P=0.2, Fig. 1D) or intramuscular fat content (P=0.7, Fig. 1E) did not differ between groups. Altogether, our results demonstrate atrophy (i.e., decrease in fiber size) and fibrotic degeneration (i.e., pathological increase in collagen content) of PFMs in parous women with symptomatic POP.

**Figure 1.**
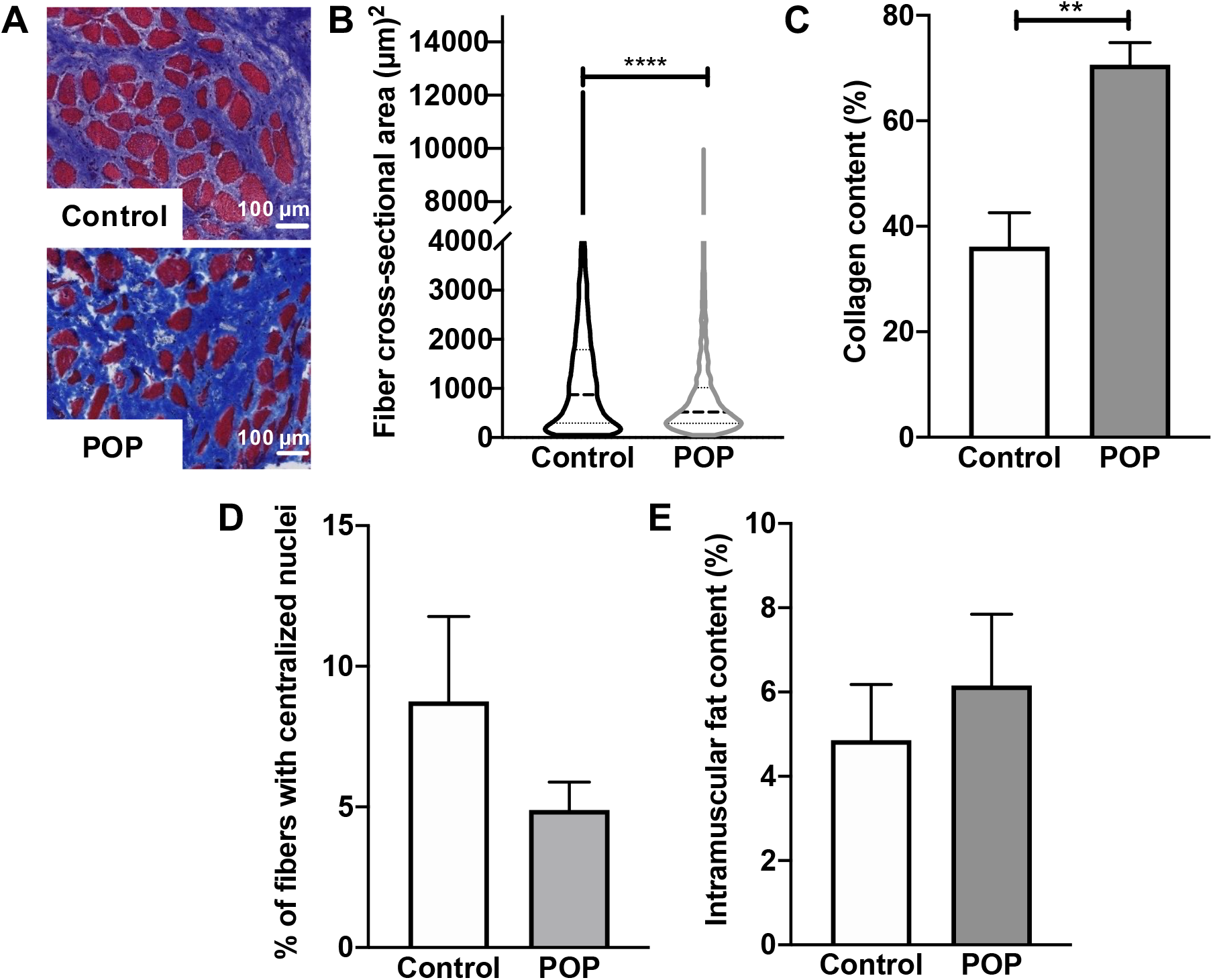
Pelvic floor muscles in women with symptomatic pelvic floor disorders demonstrate an atrophic and fibrotic phenotype. (A) Gomori’s trichrome stained cross-sections of the pubovisceralis portion of the levator ani muscle procured from vaginally nulliparous cadaveric donors without history of pelvic floor disorders (control) and parous women undergoing surgical repair of pelvic organ prolapse (POP). (B) Violin plots of fiber cross-sectional area (marker of atrophy) quantified from the trichrome images. Shape of the plots demonstrates the distribution of the myofibers of various cross-sectional areas with a median indicated by the dash line. (C) Collagen content (marker of fibrosis) quantified from Gomori’s trichrome stained muscle cross-sections. (D) Centralized nuclei quantification (marker of regeneration/degeneration). (E) Intramuscular fat content quantified from Oil-Red-O stained muscle cross-sections (not shown). N=4 (Control); N=20 (POP). *P*-values derived from Student’s t-tests for parametric and Mann-Whitney tests for non-parametric data. **p<0.01, ****p<0.0001, mean±SEM.

### Rat model of simulated birth injury replicates pathological alterations observed in parous women with pelvic organ prolapse

We previously identified that the pubocaudalis component of the rat levator ani muscle (analogous to the human pubovisceralis) undergoes the largest macro- and microscopic structural alteration acutely in response to simulated birth injury (SBI) (*13*). To assess the impact of SBI on the pubocaudalis morphological properties, we compared fiber size and collagen content at subacute (4 weeks post-SBI) and long-term (8 weeks post-SBI) timepoints. Given that revascularization has an important role in muscle regeneration (*15*), we also compared vessel density, specifically arterioles. We first compared 3 and 5-month old uninjured controls to rule out possible age-related differences between rats at the time of SBI (3-month old) and 8 weeks post-SBI (5-month old). No significant differences were observed between these groups in either the PFM fiber area (P=0.1), collagen content (P=0.3), or arteriole density (P=0.4) (Fig. S1A-C). Given the above, 3-month old uninjured controls were used to assess changes in the morphological muscle parameters consequent to birth injury.

We observed a significant decrease in PFM fiber area 4 weeks after injury compared to uninjured controls (P<0.0001). Fiber area distribution (Fig. 2A-B) revealed that injured pubocaudalis was enriched with smaller fibers. To determine whether changes in fiber size at this sub-acute timepoint were due to muscle atrophy or ongoing muscle regeneration, we assessed the centralization of nuclei, a marker of regeneration (*15*). The number of fibers with centralized nuclei was significantly increased 4 weeks post-SBI compared to controls (P=0.02), indicating ongoing regeneration (Fig. 2C). By 8 weeks post birth injury, percent centralized nuclei returned to baseline (P=0.7). Despite an increase in cross-sectional area at this longer-term timepoint compared to 4-weeks post SBI (P<0.0001, Fig. 2A-B), fiber size remained smaller than in uninjured controls (P<0.0001). The return of the centralized nuclei proportion to baseline levels with persistent smaller fiber size at 8 weeks post-SBI indicates that birth injury leads to PFM atrophy. Since myofiber atrophy is often accompanied by the pathological thickening of endo- and perimysium (*21*), we investigated whether the total amount of collagen, the major constituent of the intramuscular ECM, is altered by birth injury. We observed a significant increase in the pubocaudalis collagen content 4 weeks post-SBI compared to controls (P=0.02), which persisted at 8 weeks (P=0.03, Fig. 2D-E). These results indicate that in addition to muscle atrophy, birth injury also leads to PFM fibrosis.

**Figure 2.**
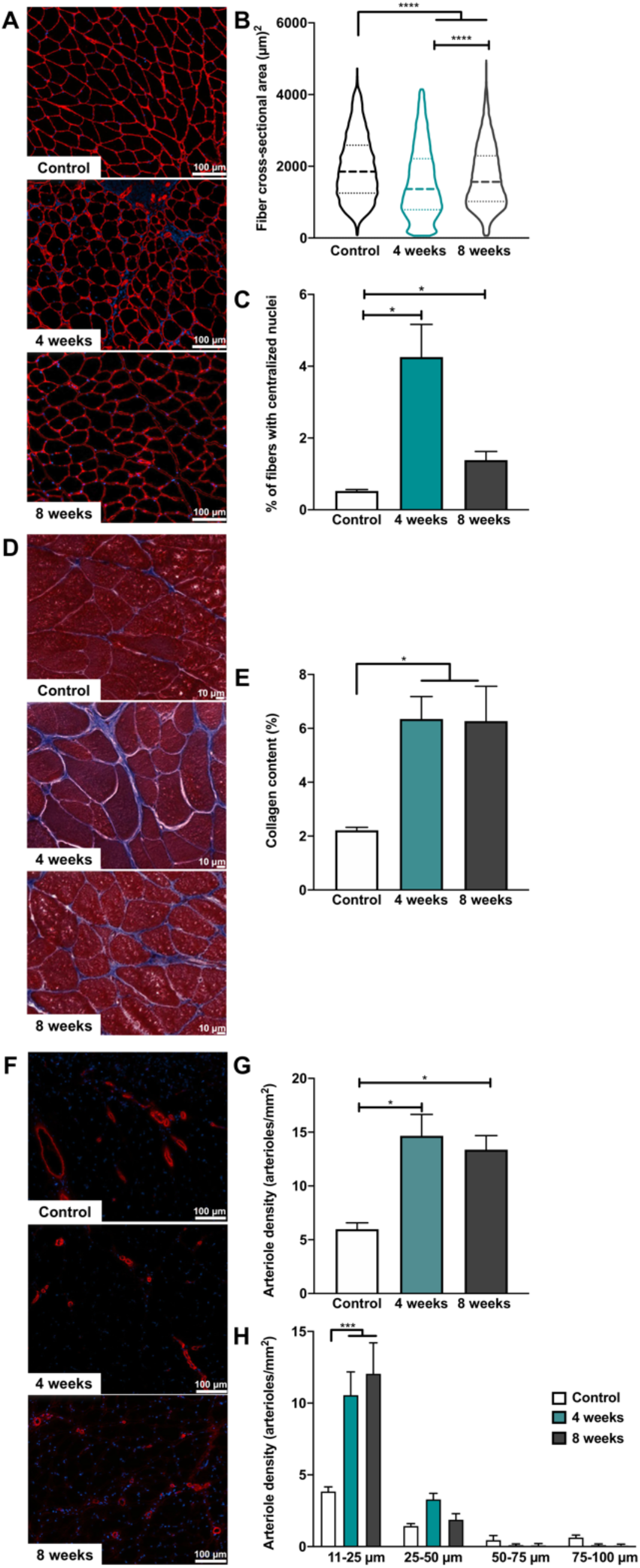
Pre-clinical rat model of simulated birth injury (SBI) recapitulating atrophic and fibrotic pelvic floor muscle phenotype observed in parous women with symptomatic pelvic floor disorders. (A) Laminin- and DAPI-stained cross-sections of the pubocaudalis portion of the levator ani muscle in uninjured controls, and 4- and 8-weeks post-SBI groups used for fiber cross-sectional area (B) and centralized nuclei (C) quantification. (D) Masson’s trichrome staining of pubocaudalis from the same groups used for intramuscular collagen quantification (E). (F) Alpha smooth muscle staining of pubocaudalis from the same groups used for quantification of the total vessel (arteriole) density (G) and distribution of vessels by size (H). N=3-6/group. *P*-values derived from one-way ANOVA followed by pairwise comparisons with Tukey’s range test for parametric data and Kruskal-Wallis followed by pairwise comparisons with Dun’s test for non-parametrically distributed data. *p<0.05, ***p<0.001, ****p<0.0001, mean±SEM.

With respect to the intramuscular vasculature, we observed an increase in total arteriole density at 4 (P=0.01) and 8 weeks (P=0.02) post-SBI relative to controls (Fig. 2F-G). We then compared the distribution of arterioles of various diameters between groups (*22*). The smaller arterioles (11-25 µm) were increased at 4 and 8 weeks post-SBI compared to controls (Fig. 2H), while no significant differences were observed in the density of larger arterioles (26-50 µm, 51-75 µm, 76-150 µm). The above indicates that birth injury alters PFM vascularization, consistent with the observations in fibrotic injured appendicular skeletal muscles (*23*).

### Myogenesis occurs within one week after simulated birth injury

To identify potential mechanisms accountable for the above long-term pathological alterations of PFMs, we elucidated, for the first time, cellular events following SBI. The unperturbed pubocaudalis demonstrated tightly packed myofibers (Fig. 3A). One day post-SBI, widespread myofiber death was observed, followed by accumulation of cellular infiltrate 3 days post injury. Regenerating myofibers were observed at 7 days, identified by centralized nuclei, with endomysial thickening at 10 days post injury (Fig. 3A). Given that muscle stem cells (MuSCs) play a key role in muscle regeneration, we next investigated their differentiation and self-renewal capacities at 1, 3, 7, and 10 days after SBI. In injured appendicular muscles, MuSCs replace or repair the damaged muscle by differentiating into myoblasts and forming new myofibers (3-4 days post-injury) or fusing with pre-existing ones, while a portion of MuSCs self-renew to replenish the stem cell pool (5-7 days post-injury) (*24, 25*). We used myogenin as a marker of differentiated MuSCs and Pax7 to identify quiescent and self-renewing cells. The peak density of differentiated MuSCs was observed 3 days post-SBI (P=0.01 vs control), with return to baseline at 7 days (Fig. 3B). The number of Pax7 expressing MuSCs increased substantially 7 days post-SBI relative to uninjured levels (P<0.0001), returning to baseline by 10 days after birth injury (Fig. 3C). The rapid differentiation of MuSCs following birth injury was confirmed by the presence of fibers positive for embryonic myosin heavy chain, a marker of de novo or fused myofibers, starting as early as 3 days post-SBI (Fig. S2). Altogether, the above data indicate that after birth injury, PFM myogenesis followed the time-course expected from the findings in the models of chemical or mechanical (*25, 26*) appendicular skeletal muscle injury. Thus, disturbance in the early regenerative process is an unlikely mechanism behind the long-term atrophic and fibrotic phenotype of the PFMs consequent to birth injury.

**Figure 3.**
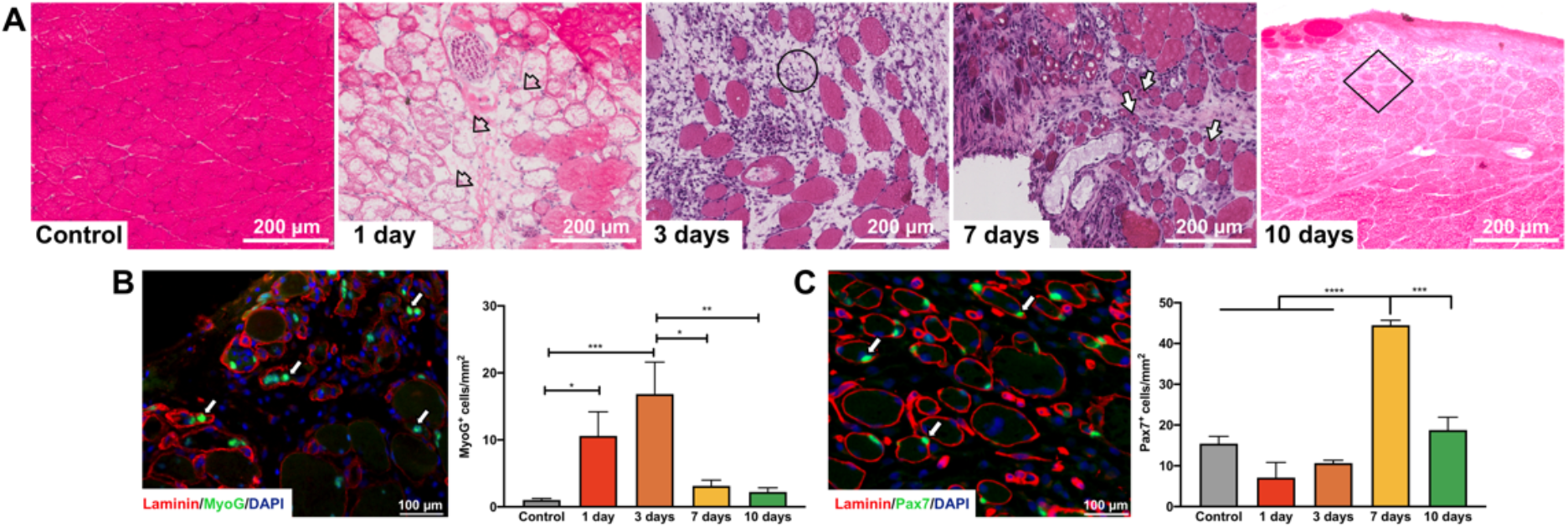
Myogenesis of the pelvic floor muscles takes place within one week after simulated birth injury (SBI). (A) Hematoxylin and Eosin staining of pubocaudalis cross-sections along a 10-day continuum following SBI. Damaged myofibers with preservation of basal lamina (arrowheads) were observed 1 day post-SBI with separation of myofibers and cellular infiltration (circle) at 3 days. Regenerated myofibers, indicated by centralized nuclei (arrows) were identified 7 days post-SBI with endomysial thickening (square) at 10 days. Muscle cross-sections were incubated with anti-Myogenin (B) and anti-Pax7 (C) antibodies for *in situ* quantification of differentiated muscle stem cells and assessment of muscle stem pool, respectively. N=3-9/group. *P*-values derived from one-way ANOVA followed by pairwise comparisons with Tukey’s range test. *p<0.05, **p<0.01, ***p<0.001, ****p<0.0001, mean±SEM.

### Sustained inflammatory response of the pelvic floor muscles follows simulated birth injury

To further decipher the mechanisms underlying the observed muscle phenotype, we next examined gene expression changes associated with PFM injury and subsequent recovery. We composed a custom panel of 150 genes from physiologically relevant pathways known to regulate injury response of the appendicular skeletal muscles. These pathways include immune response, myogenesis, muscle anabolism and catabolism, extracellular matrix (ECM) remodeling, neovascularization, and neuromuscular junction development and maturation (Table S1). We assessed the transcriptional signature of the pubocaudalis at 1, 3, 7, 10, 31, and 35 days following SBI. To determine patterns across the post-injury continuum, principal component (PC) analyses were performed for all assayed transcripts together (Fig. 4A) and separately for each individual pathway (Fig. S3A-G). Unsurprisingly, PC1 that accounts for the highest variability was attributable to the pubocaudalis response to birth injury, with progressive return towards the uninjured state during the recovery period (Fig. 4A). The same pattern was observed for each individual pathway (Fig. S3A-F).

**Figure 4.**
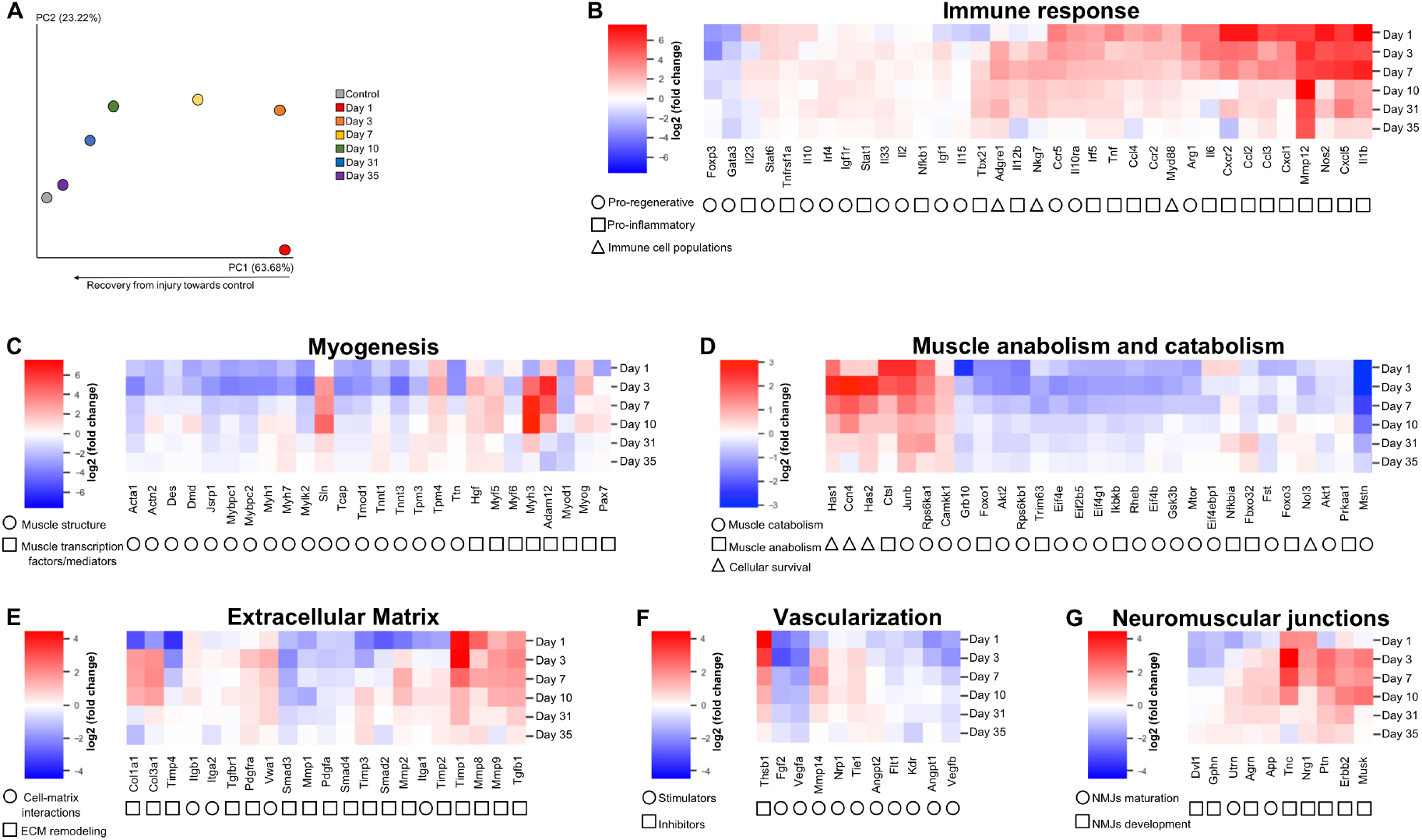
Simulated birth injury (SBI) leads to sustained inflammatory response, upregulation of ECM remodeling genes, and downregulation of genes involved in muscle anabolism. (A) Transcriptional signatures of the pelvic floor muscles across different pathways were derived from the customized Nanostring nCounter panel with 150 genes. Principal component analysis includes gene expression in uninjured controls and at multiple time points during active muscle regeneration post-SBI. (B-G) Unsupervised clustered heat maps of fold changes at each time point with respect to control. Pathways examined include immune response (B), myogenesis (C), muscle anabolism and catabolism (D), extracellular matrix (E), vascularization (F), and neuromuscular junctions (G).

We then evaluated differentially expressed genes in each pathway to determine the longitudinal alterations in gene expression in relation to the uninjured state (Tables S2-7). Figure S4 demonstrates the gene expression profile across the samples per timepoint. Clustered heat maps of the fold change with respect to uninjured controls are shown in Figure 4B-G. For the immune response pathway (Fig. 4B), pro-inflammatory genes increased dramatically following injury, with sustained upregulation of Il1b, Tbx21 (T helper cells type 1) and Irf5 (Macrophages type 1) until day 31 post-SBI. In contrast, genes associated with the pro-regenerative phase were upregulated until day 7. Notably, Foxp3, which encodes for regulatory T cells involved in the polarization of immune response towards the pro-regenerative phase, was downregulated. For myogenic pathway (Fig. 4C), expression of genes related to muscle structure predominantly decreased in response to injury, returning toward the uninjured state by 31 days. In contrast, genes coding for muscle transcription factors had variable expression. Myog, Myh3, and Pax7 expression decreased 1 day post injury. This initial decrease was followed by increased expression of Myog and Myh3 at 3 days and Pax7 at 7 days post-SBI, mimicking the differentiation/self-renewal pattern observed by immunostaining (Fig. 3B-C). For muscle anabolism pathways (Fig. 4D), we observed downregulation of all genes related to the AkT/mTOR signaling until day 10, with decreased expression of its downstream target until day 31. For muscle catabolism pathways, the majority of the genes were decreased at all time points, with the exception of Ctsl gene —involved in protein and lysosomal degradation— that increased at days 1 and 3 (Fig. 4D). For the ECM remodeling pathways (Fig. 4E), increased expression of an ECM remodeling gene, Pdgfra, was observed until day 7, while others continued upregulated until 31 days post injury, particularly Tgfb1 and Timp1. In addition, increased expression of Col1a1 and Col3a1 occurred from 3 to 10 days post-injury. For vascularization pathways, genes that promote vascularization were decreased until day 31, while genes that inhibit vascularization were increased between 1 and 10 days (Fig. 4F). Genes related to neuromuscular junction development (Nrg1, Tnc) were increased from day 1 on, while the expression of genes associated with neuromuscular junction maturation (App) started to rise at day 7 (Fig. 4G). Overall, birth injury led to impairment in PFM anabolism, sustained upregulation of ECM remodeling genes, and a persistent inflammatory response.

### Skeletal muscle ECM hydrogel (SKM) prevents pelvic floor muscle atrophy and mitigates fibrosis when administered at the time of simulated birth injury (SBI)

Given incomplete PFM recovery from SBI, we went on to investigate whether SKM, an ECM hydrogel derived from decellularized porcine skeletal muscle, could potentiate muscle recovery when injected directly into pubocaudalis at the time of SBI. We hypothesized that the immunomodulatory properties of the SKM would counteract the sustained PFM inflammatory response to SBI by inducing an early transition to the pro-regenerative environment (*20*). Injection of saline served as an experimental control (Fig. 5A). Data from animals at 4 weeks following SBI without any injection were used for comparison, given the reproducibility of our model (Fig. S5). Fiber area and collagen content were quantified 4 weeks post-SBI+injection. While saline injection increased fiber area compared to untreated SBI (P<0.0001), SKM injection resulted in a substantially greater increase in fiber area compared to saline (P<0.0001), with return of fiber size to the uninjured levels (SBI+SKM vs controls: P>0.99, Fig. 5B). SKM also lowered collagen deposition relative to untreated SBI (P=0.02), bringing intramuscular collagen content down to the level not significantly different from uninjured pubocaudalis (P=0.6). However, this response was similar to that observed in the SBI+saline group (P=0.5, Fig. 5C). These results indicate that SKM prevents PFM atrophy, restoring fiber size to the uninjured state, and mitigates fibrotic degeneration of the PFMs consequent to birth injury. With respect to vascularization, the arteriole density in the SKM group did not differ significantly compared to saline (P=0.5), untreated SBI (P=0.4), or uninjured controls (P=0.1); whereas it was significantly higher in the saline group compared to uninjured controls (P=0.01, Fig. 5D). Analysis of the arteriole density by size revealed that the smaller arterioles (11-25 µm) decreased in SKM group relative to untreated SBI (P=0.002); however, this decrease did not reach statistically significant difference compared to saline (P=0.09, Fig. 5E). The above suggests that SKM mitigates the arteriole-level vascular alterations of PFMs observed after birth injury.

**Figure 5.**
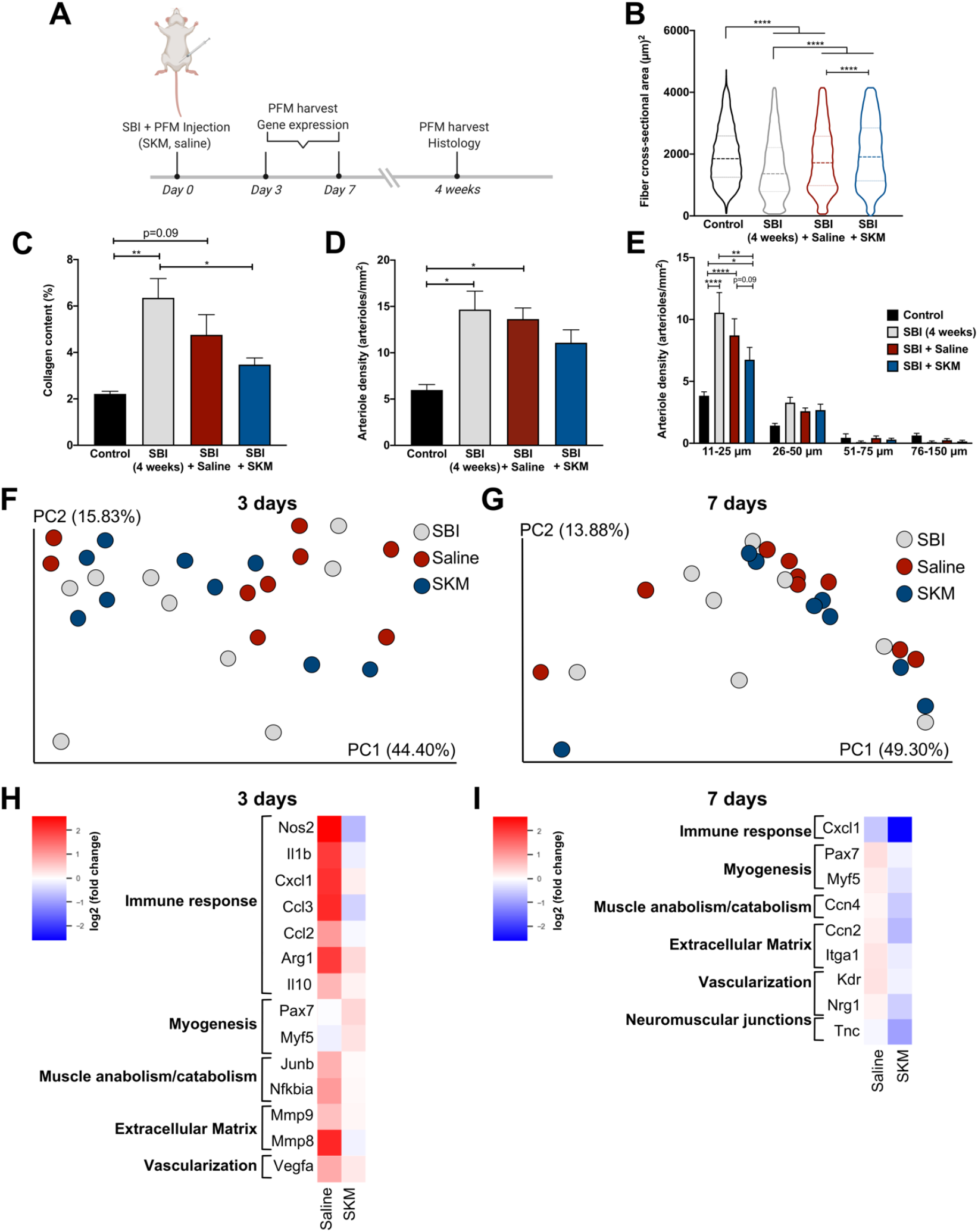
Injection of skeletal muscle extracellular matrix hydrogel (SKM) at the time of simulated birth injury (SBI) prevents pelvic floor muscle atrophy and mitigates fibrosis. (A) Study timeline. Pubocaudalis muscle of uninjured controls; and animals subjected to SBI, SBI + saline injection, or SBI + SKM injection were compared with respect to fiber cross-sectional area (B); collagen content (C); overall vessel (arteriole) density (D); and vessel (arteriole) density separated by size (E). Principal components analysis of transcriptional signatures at 3 (F) and 7 (G) days post-SBI with and without injection. Supervised heat map of fold changes for SKM and saline with respect to untreated SBI at 3 (H) and 7 (I) days. Heat map includes significantly differentially expressed genes based on the pairwise comparison between SKM and saline groups. N=6/group for histological assessments, and 8-9/group for gene expression analyses. *P*-values derived from one-way ANOVA followed by pairwise comparisons with Tukey’s range test for parametric data and Kruskal-Wallis followed by pairwise comparisons with Dun’s test for non-parametrically distributed data. Gene expression analysis was analyzed based on NanoStringDiff package in R. *p<0.05, **p<0.01, ****p<0.0001, mean±SEM.

We went on to assess transcriptional regulation of the changes in the PFM phenotype induced by SKM injection at the time of birth injury. Pubocaudalis was harvested 3 or 7 days post-SBI+injection (Fig. 5A) and analyzed with the same Nanostring panel (Table S1) for comparisons with the untreated SBI group. These timepoints correspond to the early inflammatory response and peak cell infiltration into the material, respectively. Based on the principal component analysis, SKM, saline and untreated SBI groups did not cluster separately at either time point (Fig. 5F-G); however, several genes were differentially expressed. Gene expression profiles across the samples per each timepoint are demonstrated in Figures S6-7. Clustered heat maps for each pathway, indicating fold change of SKM and saline in relation to untreated SBI and with differentially expressed genes between SKM and saline, are shown in Figures 5H-I, with pairwise comparisons illustrated in Figure S8.

At the earlier timepoint (3 days post SBI), SKM decreased the inflammatory response, as indicated by downregulation of Nos2, Il1b, Cxcl1, Ccl2, Ccl3, all of which are associated with a pro-inflammatory environment. In addition, we observed upregulation of genes associated with MuSC pool expansion (Pax7, Myf5) and a trend towards upregulation of a muscle transcription factor that encodes for de novo or fused myofibers (Myh3, P=0.06) at this timepoint. We also observed decreased expression of ECM remodeling genes at both timepoints (3 days: Mmp8, Mmp9; 7 days: Ccn2, Itga1) together with a trending downregulation of collagen I (Col1a1, P=0.07) at 7 days. Decreased expression of Vegfa and Kdr, both associated with vascularization, was observed at both timepoints. Downregulation of Tnc, involved in the development of neuromuscular junctions, was identified at 7 days. Altogether, our results indicate that compared to saline, SKM treatment at the time of birth injury decreases inflammatory response and ECM remodeling related PFM genes and upregulates genes in the myogenesis pathway early on.

### Delayed injection of skeletal muscle ECM hydrogel prevents pelvic floor muscle atrophy and mitigates fibrosis

Given that delayed injection corresponding to the timing of a routine post-partum visit may be even more clinically relevant, we next evaluated whether treatment with SKM 4 weeks post-SBI can prevent PFM atrophy and fibrosis consequent to birth injury. SKM injection 4 weeks post-SBI (Fig. 6A) potentiated muscle recovery long-term, as evident from the increased fiber area compared to saline at 8 weeks post-SBI (P<0.0001). Interestingly, SKM administration at this time point resulted in the fiber area greater than that in uninjured controls (P<0.0001) (Fig. 6B). Saline injection also increased fiber area compared to untreated SBI (P<0.0001) but failed to return fiber size to the uninjured level (P<0.0001). SKM decreased fibrotic response compared to untreated SBI (P=0.01), restoring ECM collagen content to the uninjured levels (P=0.8). Surprisingly, saline also significantly decreased collagen accumulation post injury relative to the untreated SBI group (P=0.006), resulting in the levels analogous to the uninjured controls (P=0.8), similar to the effect of SKM (P=0.9) (Fig. 6C). For arteriole density, both SKM and saline led to a decrease in the overall arteriole density (Fig. 6D), and specifically of smaller arterioles (Fig. 6E) compared to untreated SBI, restoring arteriole density to the uninjured levels (P=0.9). These results indicate that SKM administered 4 weeks after birth injury prevents PFM atrophy resulting in fiber size that supersedes that of uninjured controls and mitigates fibrosis and restore vascularization. In addition, saline injection was sufficient to also mitigate fibrosis and restore vascularization to baseline levels.

**Figure 6.**
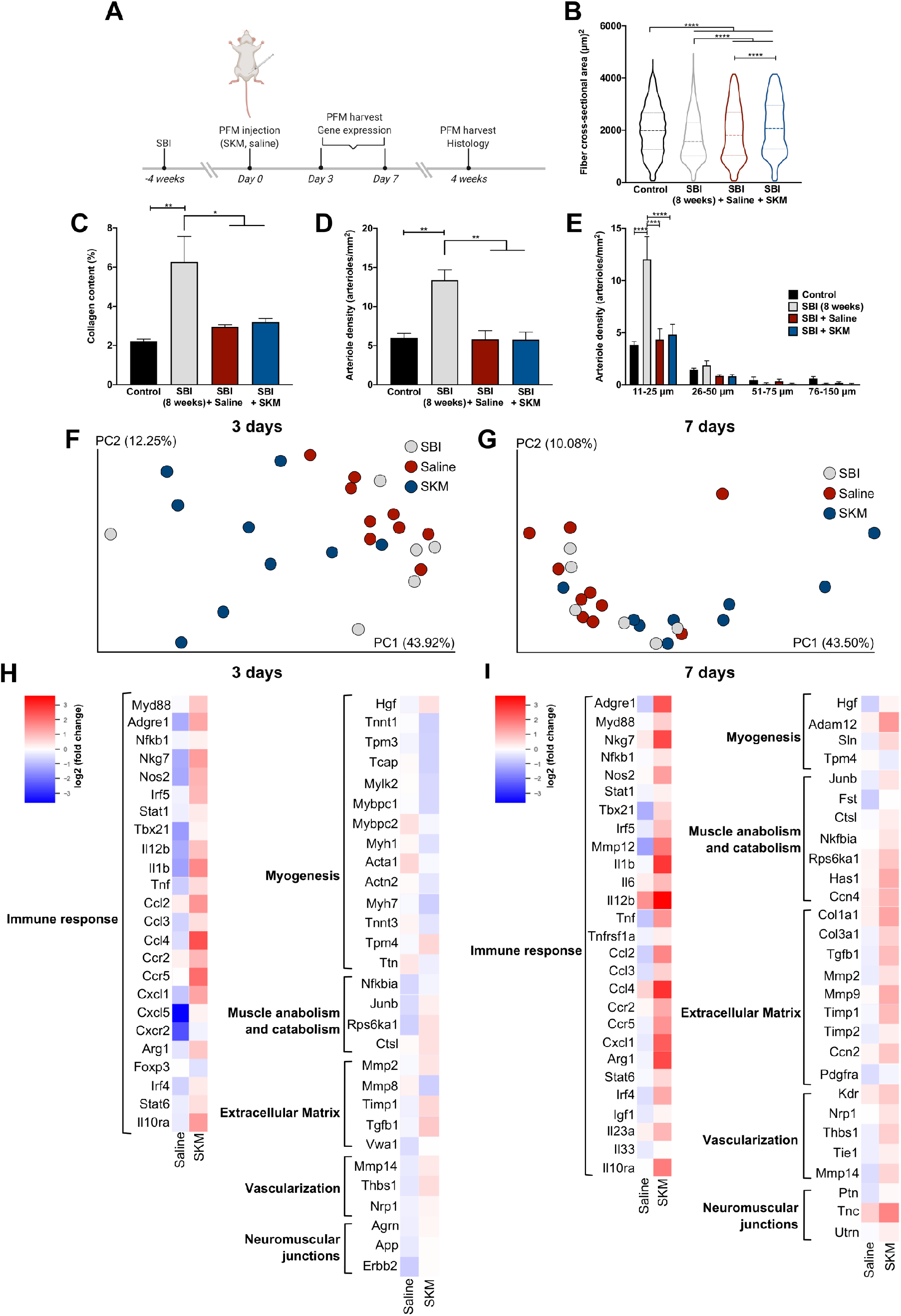
Delayed injection of skeletal muscle extracellular matrix hydrogel (SKM) prevents pelvic floor muscle atrophy and mitigates fibrosis. (A) Study timeline. Pubocaudalis muscle of uninjured controls; and animals subjected to SBI, SBI + saline injection, or SBI + SKM injection were compared with respect to fiber cross-sectional area (B); collagen content (C); overall vessel density (D); and vessel density separated by size (E). Principal components analysis of transcriptional signatures at 3 (F) and 7 (G) days post-injection (31 and 35 days post-SBI) with and without injection. Supervised heat map of fold changes for SKM and saline with respect to untreated SBI at 3 (H) and 7 (I) days. Heat map includes significantly differentially expressed genes based on the pairwise comparison between SKM and saline groups. N=6/group for histological assessments, and 6-10/group for gene expression analyses. *P*-values derived from one-way ANOVA followed by pairwise comparisons with Tukey’s range test for parametric data and Kruskal-Wallis followed by pairwise comparisons with Dun’s test for non-parametrically distributed data. Gene expression analysis was analyzed based on NanoStringDiff package in R. *p<0.05, **p<0.01, ****p<0.0001, mean±SEM.

We next analyzed the acute and subacute transcriptional changes in the PFM response to the delayed SKM injection, using the same Nanostring panel (Table S1). Pubocaudalis was harvested at 3- or 7-days post-injection, i.e. 31 and 35 days post-SBI (Fig. 6A). Despite similar response to SKM and saline with respect to intramuscular collagen content and vascularization, the principal component analysis revealed that SKM mostly clustered separately from saline and untreated SBI, especially at 3 day time point, while saline clustered together with untreated SBI (Fig. 6F-G). Gene expression profiles for each timepoint are summarized in Figures S9-10. Clustered heat maps for each pathway indicating fold change in SKM and saline groups relative to untreated SBI and with differentially expressed genes between SKM and saline are shown in Figures 6H-I. Pairwise comparisons are shown in Figure S11. At both time points, SKM led to upregulation of genes related to the immune response, specifically monocytes (Myd88), macrophages (Adgre1), macrophages type 1 (Nos2, Irf5, Stat1), macrophages type 2 (Arg1, Irf4, Stat6), T cells (Nkg7), and T helper cells type 1 (Tbx21). We identified increased expression of genes associated with pro-inflammatory (Il12b, Il1b, Tnf, Ccl2, Ccl3, Ccl4, Ccr2, Ccr5, Cxcl1, Cxcl5, Cxcr2), as well as pro-regenerative (Il10ra, Il33, Igf1) mediators. In addition, genes that code for mediators of myogenesis (3 days: Hgf; 7 days: Hgf, Adam12) and cellular survival (7 days: Has1, Ccn4) were upregulated. Genes associated with muscle structure were downregulated (3 days: Tnnt1, Tpm3, Tcap, Mylk2, Mybpc1, Mybpc2, Myh1, Acta1, Actn2, Myh7, Tnnt3, Tpm4, Ttn). While upstream genes of muscle anabolism pathway (Junb, Rps6ka1, Fst) were upregulated at 3 and 7 days, the downstream targets remained unchanged. Upregulation of ECM remodeling genes was identified at both time points (Mmp2, Mmp9, Timp1, Tgfb1, Ccn2), with increased expression of collagens I and III (Col1a1, Col3a1) at 7 days. Although receptor genes related to vascularization pathway were upregulated (Kdr, Nrp1, Tie1), the ligands were not differentially expressed. Lastly, we observed upregulation of genes involved in the development (Agrn, Erbb2, Ptn, Tnc) and maturation (App, Utrn) of neuromuscular junctions at 3 and 7 days. Taken together, at the timepoints studied, SKM increases the expression of genes associated with both pro-inflammatory and pro-regenerative immune responses, ECM remodeling, myogenesis, and neuromuscular junction development and maturation.

## Discussion

We demonstrate, for the first time, that pubovisceralis, which sustains the highest strains during vaginal delivery, exhibits substantial myofiber atrophy and severe fibrosis in parous women with symptomatic pelvic organ prolapse. The widespread degeneration observed in this group was not evident in PFMs procured from age-comparable nulliparous cadaveric donors without history of pelvic floor disorders. Furthermore, we recapitulated these pathological alterations in the rat model of simulated birth injury (SBI). Analyses of the PFM transcriptional signatures across multiple time points post SBI demonstrate a sustained inflammatory response, impairment in muscle anabolism, and persistent expression of ECM remodeling genes. These transcriptional alterations point towards putative mechanisms that account for PFM atrophy and fibrosis. We further demonstrate that an acellular injectable skeletal muscle ECM hydrogel (SKM), administered at two clinically-relevant time points, can either prevent or mitigate these untoward PFMs’ alterations caused by birth injury.

Maternal PFM birth injury and subsequent muscle dysfunction are major inciting events that lead to the development of pelvic floor disorders, especially pelvic organ prolapse, later in life. Radiologically detected PFM avulsions have been the widely acknowledged phenotype associated with pelvic floor disorders and the primary focus of the literature to date. However, the radiological evidence of such mechanical uncoupling is not present in all women with decreased PFM strength, suggesting the existence of other causes of PFM dysfunction. Furthermore, published clinical studies indicate that a sizable proportion of PFM avulsions observed on imaging shortly after childbirth are no longer evident 1 year postpartum (*11*), while large-scale cadaveric investigations demonstrate absence of radiologically-detected “avulsions” on direct anatomic dissections (*10*). Together with our findings in the rat model (*13*), the above supports our notion that PFM birth injury can result in pathological transformations, other than avulsions, not detectable by routine imaging modalities. Expanding our understanding of PFM injuries beyond mechanical uncoupling is of utmost importance to foster identification of novel therapeutic targets and approaches. Previous studies have demonstrated that the morphological PFM changes associated with pelvic organ prolapse are muscle thinning and decreased volume identified with magnetic resonance imaging, indicative of muscle atrophy (*27, 28*). In addition to myofiber atrophy, we identified dramatic fibrotic degeneration of PFMs in parous women with pelvic organ prolapse. To understand the mechanism of injury leading to such phenotype, we relied on an animal model to investigate PFM response to strains associated with birth injury.

The constructive regeneration of acutely injured limb muscles that returns them to the pre-injury state is mainly completed within a 4-week recovery period (*15, 29*). In contrast, we demonstrate that the endogenous regenerative potential of PFMs is insufficient to restore the muscle to its baseline following SBI, resulting in PFM atrophy and fibrosis. Multiple studies conducted in the appendicular and trunk muscles conclude that acute skeletal muscle injury is followed by a temporally regulated immune response. The pro-inflammatory phase lasts up to 3-4 days post-injury, followed by the pro-regenerative phase that promotes tissue remodeling and repair. The transition towards the latter phase is influenced by a subpopulation of T-helper cells (Tregs) (*30*). In our study, the pro-inflammatory and pro-regenerative phases overlapped during the acute period post-SBI (Fig. S12) with a sustained inflammatory response until day 31 post-injury, which could be attributed to the downregulation of Tregs. This cellular population is recruited after muscle injury by the Il33 cytokine. Kuswanto, et al., demonstrated that reduced Il33 expression acutely after muscle injury leads to an impairment in Treg accumulation (*31*). Even though the increased expression of Il33 was present at 3 days after injury in our study, we opine that this expression was insufficient to influence the recruitment of Tregs. Our data suggest that the chronic inflammatory response of PFMs following SBI negatively impacts pathways related to muscle growth, acute ECM remodeling, and vascularization.

Skeletal muscle growth and repair occur via myogenesis (i.e., cellular turnover) or through increase in protein synthesis (i.e, anabolism). In limb muscles, the source of de novo myofibers observed acutely after muscle injury is committed MuSCs poised for immediate differentiation without proliferation. Following this swift differentiation, between 5 to 7 days post injury, the majority of remaining MuSCs self-renew to replenish the stem cell pool (*24*). In our study, the early brief upregulation of pro-regenerative mediators correlated with the increased expression of genes associated with the MuSC differentiation and myofiber regeneration from 1 to 10 days (Fig. S12) and the increased density of differentiated Myog^+^ MuSCs at 3 days post SBI (Fig. 3B). Consistently, the upregulation of pro-inflammatory cytokines coincided with the upregulation of genes involved in MuSC expansion from 3 to 10 days (Fig. S12); this was further correlated with the increased density of Pax7^+^ MuSCs at 7 days (Fig. 3C). Given that myogenesis of pubocaudalis followed the expected time course after SBI, we focused on the protein turnover pathways to determine potential mechanism responsible for PFM atrophy. The AkT/mTOR pathway plays a key role in protein synthesis and its activation is negatively influenced by prolonged inflammation (*32*). Our data suggest that persistent inflammation after birth injury could be a culprit in the downregulation of the AkT/mTOR pathway, as indicated by the persistent downregulation of one of its main downstream targets (Rpskb1) until 31 days post injury. Furthermore, Il1, known to promote decrease in mTOR signaling, leading to muscle atrophy (*32*), was upregulated until day 31. The principal mechanism for activation of the AkT/mTOR pathway involves phosphorylation of its components, which we did not assess in the current study; however, published studies demonstrate correlation of this signaling pathway at the transcriptional, translational and post-translational levels during muscle anabolism (*33, 34*).

Following acute muscle injury, a temporal increase in ECM deposition is crucial to provide a scaffold that regulates MuSC function and muscle repair. However, excessive ECM production and dysregulation of ECM metabolism leads to a pathological increase in intramuscular ECM components, mainly collagen I (*21*). ECM remodeling mediators are secreted by damaged myofibers, immune cells, and fibroadipogenic progenitors (FAPs), which also support MuSC differentiation (*35*). In our study, upregulation of Pdgfra, a gene that encodes the marker of FAPs in skeletal muscles (*35*), was followed by the increase in genes related to MuSC differentiation and ECM production from 3 to 10 days (Fig. S12). Although we observed an expected upregulation of genes associated with the temporal ECM deposition, we suggest that the sustained expression of Tgfb1, a master regulator of fibrosis, and Timp1, involved in inhibition of ECM degradation (*21*), lead to the long-term pathological increase in PFM collagen content.

After muscle injury, inflammation has been shown to alter vascularization with unknown overall impact on the intramuscular blood flow (*15, 23*). In our study, the sustained inflammatory response was associated with the increased vessel number, specifically smaller arterioles. At the gene expression level, we observed upregulation of the indirect stimulators of arteriogenesis (Ccl2, Tnfa, Tgfb1); however, direct stimulators (Fgf2, Vegfa, Vegfb, Flt1, Kdr, Angpt2) (*36*) were downregulated. Due to our targeted biased approach, we did not investigate other direct stimulators (Fgf1, Fgf4 or Pdgfb) that could also contribute to arteriogenesis.

After characterizing PFM *in vivo* response to birth injury, we focused on exploring a regenerative strategy to potentiate constructive muscle remodeling after SBI. Current preventative and therapeutic approaches for PFM dysfunction are limited to PFM rehabilitation, plagued by low overall success rate and poor long-term adherence (*16*). Among regenerative strategies, cell-based therapies introduce challenges related to survival and engraftment, potential tumorigenicity, high costs and short shelf life (*37, 38*). On the other hand, acellular immunomodulatory biomaterials can be used to potentiate endogenous regeneration (*39*), as we have previously shown in ischemic limb muscles (*20*). Here, we investigated whether SKM could be used to treat injured PFMs at the time of SBI or 4 weeks post injury. Our overarching goal is to develop therapeutic approaches for vaginally parous women - the most vulnerable population with respect to PFM injury and pelvic floor disorders. Thus, these time points were chosen to reflect two translationally relevant opportunities to use SKM. As a preventative measure, SKM could be used on Labor and Delivery for women at high risk for PFM injury (i.e. operative vaginal delivery, prolonged second stage of labor, macrosomia, obstetrical laceration) (*40*). As a therapeutic intervention, SKM could be used for women with clinically detected PFM weakness, identified by digital palpation or with the use of perineometer (*41*) at the post-partum visit, routinely scheduled within 2 months after childbirth. Moreover, SKM could be injected with minimal invasiveness directly into PFMs, identified by palpation, a technique familiar to most obstetricians and gynecologists, or with an ultrasound available on Labor and Delivery units and in outpatient office settings. A similar injectable ECM hydrogel derived from porcine myocardium was developed and tested in a Phase I clinical trial in myocardial infarction patients (*42, 43*), which demonstrates the translational potential of an ECM hydrogel therapy.

Immediate delivery of SKM following SBI led to decreased expression of several key pro-inflammatory genes, although pro-regenerative genes were not impacted at the time points studied. With respect to the observed improvement in PFM phenotype, we suggest that the increase in PFM fiber area in response to SKM is due to the material’s enhancement of the expanding MuSC population rather than upregulation of the muscle anabolism, as the downstream targets of the mTOR pathway were unchanged. Indeed, upregulation of genes associated with MuSCs together with a trending increase of genes related to de novo or fused myofibers were observed early on (3 days). Previous studies, indicating that SKM increases the MuSC recruitment (*20, 44*), are consistent with our current findings that SKM supports PFMs’ stem cell function. With respect to the intramuscular collagen content, we suggest that SKM mitigates fibrosis by downregulating ECM remodeling genes, such as Ccn2 (connective tissue growth factor), involved in ECM deposition (*21*). Lastly, immediate injection of SKM mitigates the altered vascularization, observed at the levels of arterioles, after birth injury. The above could be an indirect consequence of the downregulation of pro-inflammatory related genes or a direct result of decreased Vegfa expression (*36*).

Interestingly, although the long-term improvements in PFM phenotype were similar after SKM administration at either time point, the differential gene expression in response to SKM vs saline was more prominent for the delayed injection (Fig. 6F, G). One month after birth injury, the tissue environment is not as dynamic as immediately after SBI. ECM hydrogels are known immunomodulators (*45*), and indeed we found shifts in both the pro-inflammatory and pro-regenerative responses. While delayed SKM injection did not upregulate genes directly associated with MuSC function, it upregulated genes that aid in myogenesis, specifically Hgf and Adam12. The former is involved in the migration and activation of MuSCs (*46*), while the latter one is expressed transiently during muscle regeneration at the time of MuSC differentiation and fusion (*47*). Thus, similar to the effects of SKM at the time of birth injury, biomaterial administration 4 weeks post SBI appears to increase the PFM fiber size via SKM impact on the myogenesis, rather than muscle anabolism. In addition, both SKM and saline mitigated fibrosis and restored arteriole density to the uninjured levels, which is, at least partially, a likely consequence of the acute inflammatory response to the micro-injury associated with the process of injection, although the gene expression profile following SKM or saline injection did differ. Lastly, delayed injection of SKM potentiated expression of genes associated with neuromuscular junction development and maturation, which contribute to overall muscle repair.

One of the limitations of our study relies on a biased approach in designing the custom gene panel. We selected key genes based on the published literature to cover different pathways relevant to skeletal muscle regeneration, however, we are aware that this limits an unbiased and in-depth analysis of pathways’ regulation following PFM injury and biomaterial injection. Secondly, we performed our studies in non-pregnant model to obviate the potential confounding effects of the complex hormonal milieu associated with pregnancy, size and number of pups, and the effects of spontaneous parturition.

In conclusion, our study demonstrates that PFMs in parous women with symptomatic pelvic floor disorders exhibit severe atrophy and fibrotic degeneration. In addition, we show that the rat birth injury model recapitulates this phenotype, with long-term muscle atrophy and fibrosis associated with sustained inflammatory response. Finally, we provide evidence to support the novel use of a minimally-invasive acellular regenerative biomaterial therapy at two clinically relevant interventional time points to prevent and mitigate pathological alterations of PFMs consequent to birth injury.

## Materials and Methods

### Study Design

The objectives of this study were to: 1) determine and compare morphological properties of PFMs in parous women with symptomatic pelvic organ prolapse and age-comparable nulliparous cadaveric donors without history of pelvic floor disorders; 2) investigate the endogenous response of PFMs to birth injury along the biologically-relevant continuum using validated pre-clinical model of simulated birth injury (SBI); and 3) investigate the efficacy of skeletal muscle extracellular matrix hydrogel (SKM) at two translationally relevant time points following birth injury. Animals were randomized for the biomaterial-related studies. The sample size, indicated in the figure legends, was calculated *a priori* (see Statistical Analysis). No outliers were excluded from the study. For the quantitative analyses, the investigators were blinded to the group identity.

### Pelvic Floor Muscle Collection from Women with Pelvic Organ Prolapse

The pubovisceralis biopsies were obtained during surgical pelvic organ prolapse correction (n=20). Study participants were enrolled in the Institutional Review Board approved study at two sites— Kaiser Permanente San Diego and the University of California San Diego. Relevant obstetrical, surgical, and medical history were collected for each subject. Immediately after procurement, muscle biopsies were pinned to a cork at slack length, snap-frozen in isopentane chilled with liquid nitrogen, transported on dry ice, and stored in −80°C. At the time of tissue processing, tissue was embedded in optimal cutting temperature material for cryosectioning and snap-frozen again. Ten micron-thick cross-sections were stained with Hematoxylin and Eosin for myofibrillar shape and packing, Oil-red-O for intramuscular fat content, and Gomori’s trichrome for collagen content, fiber area, and centralized nuclei, using well-established methods.

### Cadaveric Pelvic Floor Muscle Collection

Nulliparous cadaveric donors without history of pelvic floor disorders were used as controls (n=4). Samples were obtained through the Bequest Body Donation Program at the University of Minnesota, which provides relevant obstetrical, surgical, and medical history for each donor. Donors with history of gynecologic or colorectal malignancy, pelvic metastasis, pelvic radiation, connective tissue disorder, myopathy, rectal prolapse, colectomy, or proctectomy were excluded. Biopsies of pubovisceralis were obtained from the muscle mid-belly within 7 days post-mortem and immediately processed as described above.

### Simulated Birth Injury Rat Model

All procedures were approved by the Institutional Animal Care and Use Committee at the University of California, San Diego. Three-month-old Sprague-Dawley female rats (Envigo, Indianapolis, IN) were anesthetized using 2.5% isoflurane with oxygen for the duration of the procedure. Vaginal distention was performed using an established protocol (*13*). Briefly, a 12-French transurethral catheter (Bard Medical, Covington, GA) with the tip cut off was inserted into the vagina with 130 grams weight attached to the end of the catheter. The balloon was inflated to 5 ml and left in place for 2 hours, after which it was pulled through the introitus to replicate circumferential and downward strains associated with parturition.

### Rat Pelvic Floor Muscle Collection to Assess Endogenous Response to Simulated Birth Injury

For subacute and long-term outcomes, animals were sacrificed and pubocaudalis was harvested 4 or 8 weeks post-SBI. To assess early response to SBI, pubocaudalis was harvested at 1, 3, 7, or 10 days post-injury to assess the overall tissue morphology by H&E and to quantify phases of myogenesis by immunohistochemistry. Harvested muscles were embedded in optimal cutting temperature material and snap-frozen in isopentane chilled with liquid nitrogen. Expression of genes relevant to muscle regeneration was performed using Nanostring to investigate putative mechanisms accountable for the long-term muscle phenotype. Pubocaudalis was harvested at 1, 3, 7, 10, 31 and 35 days post-SBI—time points of ongoing muscle regeneration. Muscles were submerged in RNAlater™, stored at 4°C overnight, and transferred to −80°C before RNA isolation.

### Pelvic Floor Muscle Collection after Biomaterial Injection

SKM was fabricated and characterized as described in the supplementary materials (Fig. S13, Table S8). For the immediate injection, animals were subjected to SBI, followed by 10 µl saline (experimental control) or SKM injection directly into the rostral portion of pubocaudalis. For the delayed injection, either saline or SKM (10 µl) was injected 4 weeks post-SBI. The minimally invasive injection via transobturator approach (Fig. S14A) was performed using Hamilton syringe with a 30G needle. To assure reliable SKM injection, we first performed multiple injections with biomaterial prelabeled with India ink or Alexa fluor 568 (Fig. S14B-C). The material was consistently observed in the rostral/enthesial portion of pubocaudalis. This region of pubovisceralis and pubocaudalis is known to experience the highest strains during the human parturition and SBI in the rat model, respectively (*9, 13*). We performed a long-term efficacy study, where saline and SKM groups were compared to each other and to the untreated SBI group. We then performed another study to assess gene expression changes at 3 and 7 days post-injection. Methods for histological analysis, RNA isolation, and Nanostring gene expression analysis are included in supplementary materials.

### Statistical Analysis

Sample size for histological analysis was determined based on the previous study assessing efficacy of SKM (*20*). Using G*power software, 6 animals/group/timepoint were needed to achieve 90% power at a significance level of 0.05. For gene expression studies, preliminary data with qRT-PCR were used to obtain sample size. Ten animals/group/timepoint were needed to achieve 80% power at a significance level set to 0.05. Data that followed a parametric distribution were compared using a Student’s t-test, or a one-way analysis of variance followed by Tukey’s post hoc pairwise comparisons. Non-parametrically distributed variables, such as fiber area, were analyzed by Mann-Whitney or Kruskal-Wallis test followed by Dunn’s pairwise comparisons (*48, 49*). Data were analyzed using GraphPad Prism v8.0, San Diego, CA. Gene expression normalization and differential expression was analyzed using the NanoStringDiff package in R, with a significance at a p value<0.05 and a fold change cutoff of 1±0.25 (*50*).

## Acknowledgments

The authors thank the individuals who donated their bodies to the University of Minnesota’s Anatomy Bequest program for the advancement of education and research.

## Author contributions

PD contributed to the design of the experiments, performed in vivo experiments, processing of biomaterial, tissue collection and processing, image analysis, gene expression panel design and analysis, and wrote the manuscript. FB contributed to the experimental design and conduct, and data interpretation related to muscle stem cells and muscle regeneration, as well as manuscript writing and editing. LB performed data visualization for gene expression studies and to the manuscript writing and editing. MC enabled collaboration with the Bequest Body Donation program and collected all cadaveric specimens. GZ and SM contributed to the enrolment and consenting of the study subjects, and tissue procurement from living women. ED, contributed to tissue processing and imaging analysis, SF and MMS contributed to imaging analysis. CD and MS wrote algorithms for quantitative imaging analysis of human samples. KLC contributed to the experimental design, development of biomaterial, project coordination, data interpretation, and manuscript editing. MA contributed to the experimental design, overall project coordination, human and animal specimen collection, in vivo animal experiments, data interpretation, and writing of the manuscript.

## Data and materials availability

All data associated with this study are available in the main text or the supplementary materials.

## Supplementary materials

### Materials and Methods

Figure S1. Pelvic floor muscle fiber cross-sectional area, total collagen content, and overall arteriole density do not differ between 3-vs 5-month old rats.

Figure S2. Representative images of regenerating myofibers after simulated birth injury.

Figure S3. Principal component analysis of transcriptional signatures during active regeneration of the pelvic floor muscles following simulated birth injury.

Figure S4. Gene expression profile across the samples per timepoint post-simulated birth injury.

Figure S5. Reproducibility of simulated birth injury (SBI).

Figure S6. Gene expression profile across the samples 3 days after immediate injection of skeletal muscle extracellular matrix (SKM) hydrogel.

Figure S7. Gene expression profile across the samples 7 days after immediate injection of skeletal muscle extracellular matrix (SKM) hydrogel.

Figure S8. Pairwise comparisons after immediate injection of skeletal muscle extracellular matrix (SKM) hydrogel.

Figure S9. Gene expression profile across the samples 3 days after delayed injection of skeletal muscle extracellular matrix (SKM) hydrogel.

Figure S10. Gene expression profile across the samples 7 days after delayed injection of skeletal muscle extracellular matrix (SKM) hydrogel.

Figure S11. Pairwise comparisons after delayed injection of skeletal muscle extracellular matrix (SKM) hydrogel.

Figure S12. Schematic representation of the interplay between different pathways analyzed at multiple time points following simulated birth injury.

Figure S13. Characterization of skeletal muscle extracellular matrix (SKM).

Figure S14. Intramuscular injection of decellularized skeletal muscle extracellular matrix (SKM) hydrogel.

Table S1. Customized Nanostring nCounter panel list.

Table S2. Immune response related genes throughout the different time points after simulated birth injury.

Table S3. Myogenesis related genes throughout the different time points after simulated birth injury.

Table S4. Muscle anabolism and catabolism related genes throughout the different time points after simulated birth injury.

Table S5. Extracellular matrix remodeling related genes throughout the different time points after simulated birth injury.

Table S6. Vascularization related genes throughout the different time points after simulated birth injury.

Table S7. Neuromuscular junctions related genes throughout the different time points after simulated birth injury.

Table S8. Composition of decellularized porcine skeletal muscle ECM.

