## Supplementary Materials for "Pro-regenerative Extracellular Matrix Hydrogel Prevents and Mitigates Pathological Alterations of Pelvic Muscles Following Birth Injury"

Materials and Methods

*Histological analysis*

Hematoxylin and Eosin (H&E), Oil-red-O, Masson's Trichrome (Polysciences kit) stainings were done as described previously (*51*). Immunohistochemistry was done to stain laminin, vessels, quiescent and activated muscle stem cells, and differentiated muscle stem cells. Histological analyses were done with a Leica Aperio ScanScope® CS$^2$ (H&E), Leica Ariol® (Laminin, vessels), and Leica DM600B (muscle stem cells). For fiber area and centralized nuclei, anti-laminin antibody (Abcam, Cambridge, MA; 1:200) was used and an Alexa Fluor 488 conjugated secondary antibody (Invitrogen, Carlsbad, California; 1:500), and nuclei were counterstained with Hoechst 33342. Fiber area was measured using a custom ImageJ (NIH, Bethesda, MD) Macro for a total of approximately 10,000 fibers per sample, covering 8 tissue sections. Fibers with centralized nuclei were manually counted from the same tissue sections and divided by the total number of fibers per section. For vessels staining, arterioles were stained with an alpha-smooth muscle actin (αSMA) antibody (Dako, Carpinteria, California; 1:75) and an Alexa Fluor 488 conjugated secondary antibody (Invitrogen, Carlsbad, California; 1:500), and nuclei were counterstained with Hoechst 33342. Five regions of interest per tissue section at 10X were obtained throughout 5 sections per sample. The midline of arterioles was traced and the average feret (maximum and minimum) diameter was recorded using ImageJ. Muscle stem cells density was assessed by an anti-Pax-7 antibody (DSHB, Iowa city, Iowa; 1:100), co-stained with anti-laminin antibody (Abcam, Cambridge, MA; 1:200) and an Alexa Fluor 488 and 594 conjugated antibodies (Invitrogen, Carlsbad, California; 1:250), respectively, and nuclei were counterstained with Hoechst 33342. Pax-7$^+$ cells located below laminin were quantified in 4 regions of interest per section at 20X, for a total of 8 sections per sample. Number of cells was normalized to area per image, to obtain cell density. Differentiated muscle stem cells was performed with anti-myogenin antibody (Abcam, Cambridge, MA; 1:200), co-stained with anti-laminin antibody (Abcam, Cambridge, MA; 1:200) and an Alexa Fluor 488 and 594 conjugated antibodies (Invitrogen, Carlsbad, California; 1:250), respectively, and nuclei were counterstained with Hoechst 33342. Imaging analysis was performed as described for Pax7.

*RNA isolation and Nanostring*

Samples were thawed, homogenized with a sonicator (TissueRuptorII, Qiagen, Germantown, Maryland) and RNA was isolated with RNAeasy Fibrous Tissue Mini Kit following manufacturer instructions (Qiagen, Germantown, Maryland). We used NanoString nCounter ® MAX Analysis System with an nCounter® custom codeset designed for rat with different categories take into consideration (Table S1 for a complete gene list). Briefly, RNA concentration was measured using a Qubit 3.0 Fluorometer with a Qubit™ RNA HS Assay kit. The hybridization buffer (70 µl) was then mixed with the Custom Reporter CodeSet solution, and 8 µl of this master mix was then added separately to 50-100 ng of RNA per tissue sample, and RNA-free water up to 13 µl total. Then, 2 µl of Capture ProbeSet was added to the mixture, thoroughly mixed and placed on a thermocycler at 65°C for 16-48 hours and then maintained at 4°C for less than 24 hours. Using a two-step magnetic beads purification, probe excess was removed in PrepStation and target/probe complexes were bound on the cartridge. The data was collected by the digital analyzer (NanoString nCounter® Digital Analyzer) with images of immobilized fluorescent reporters in the sample cartridge. Results of barcode reads were analyzed by nSolver™ Analysis Software 4.0 and differential expression analysis was done with a custom R script. The nanostring data were visualized using QIIME 2 (https://view.qiime2.org/) and ggplot and pheatmap packages in R.

*Skeletal muscle ECM hydrogel fabrication*

The process for development of the porcine skeletal muscle ECM was done as previously described (*20, 52, 53*). Briefly, longissimus dorsi muscle was harvested from Yorkshire farm pigs (30-45 kg), immediately after euthanasia. Subcutaneous fat and connective tissue were removed. The skeletal muscle was then cut into small pieces of 3-5 mm in length, placed in beakers of approximately 30 grams per beaker and rinsed in water for 30-45 min. Then, the tissue was spun in 1% (w/v) sodium dodecyl sulfate (SDS) (Fischer Scientific, Fair Lawn, NJ) in phosphate buffered saline (PBS) with 0.5% penicillin streptomycin (PS) of 10,000 U/mL (Gibco, Life Technologies, Grand Island, NY) for 2 hours. The tissue was then rinsed and daily changes of SDS solution with PS were done for 3-5 days to remove the cellular content. After the tissue was fully white, it was spun in ultrapure water for 2 hours, followed by isopropyl alcohol (IPA) (Fischer Scientific, Fair Lawn, NJ) for 18-24 hours for lipid removal. Finally, the tissue was rinsed in water for 24 hours followed by approximately 5 additional rinses in water, freezing at -80 °C, and lyophilizing (for 48 hrs). The lyophilized material was then milled into a fine powder using a Wiley® Mini-Mill and a #60 sieve. Samples from five different pigs were combined to form a batched material that was used for all experiments. The powder was then digested at 10 mg/mL in a solution of pepsin at 1 mg/mL (Sigma, St. Louis, MO) in 0.1 M HCl for 48 hours (all solutions were sterile filtered). After digestion, the liquid form was brought to a pH of 7.4 while on ice, stirring and using chilled 1.0 M NaOH. The salt concentration was adjusted to 1x PBS by adding 10x PBS. Next, the ECM concentration was lowered to 6 mg of ECM/ml with 1x PBS. Finally, the material was then frozen and lyophilized for long-term storage at -80 °C. Before using it, the material was thawed and brought back to a concentration of 6 mg of ECM/mL with sterile ultrapure water. The characterization process of the material is described on Fig. S14 and Table S8.

*Characterization of decellularized skeletal muscle ECM*

Fresh and decellularized samples were cryosectioned and staind with Hematoxylin and Eosin (H&E). Images were obtained with Leica Aperio ScanScope ® CS$^2$. DNA content was measured by isolating dsDNA (n=3) using a NucleoSpin® Tissue kit (Macherey-Nagel, Duren, Germany). The nuclei acid composition was quantified with a Thermo Scientific NanoDrop 2000c spectrophotometer and the dsDNA was measured with the Quint-iT$^{TM}$ PicoGreen® dsDNA Assay Kit (Invitrogen, Eugene, OR). A DMMB protocol (*52*) was used to measure the sulfated glycosaminoglycan (sGAG) content (n=3). SDS-PAGE was used to characterized protein fragment molecular weights (n=3).

For quantification of protein composition, ~1mg of decellularized ECM was prepared for liquid chromatography with tandem mass spectrometry (LC-MS/MS), as previously published (*53*). Briefly, material was digested for 24 hrs at room temperature (rt) in 500 µL/mg of 100 mM CNBr in 86% TFA. After digestion and water rinses, samples were neutralized with 0.1 M ammonium bicarbonate buffer (ABC), followed by another digestion with a 10 kDA molecular weight cutoff filter based on the FASP protocol (*54*). Prior to digestion, 1 pmol of $^{13}C_6$ Quantitative Concatamer (QconCAT) peptides, were added which represent ECM and ECM-associated proteins (*55*). Samples were rinsed with 8 M urea and 0.1 M ABC pH 8.5, followed by 10 mM DTT for 30 minutes at rt, and alkylated with 55 mM iodoacetamide for 30 minutes at rt. Then, samples were centrifuged, washed with urea and 10 mM ABC pH 8 for 3 times. Proteins were digested at 37°C overnight with modified trypsin (Promega, Madison, WI) at a concentration of 1/50 protease/protein (wt/wt) in 0.02% ProteaseMax surfactant (Promega, Madison, WI). Peptides were eluted with 10 mM ABC pH 8.0, dried on a Speed-Vac (ThermoFisher Scientific, Waltham, MA), and reconstituted with a known volume of control alcohol dehydrogenase peptide for quantitative normalization. For liquid chromatography-selected reaction monitoring (LC-SRM), a QTRAP® 5500 triple quadrupole mass spectrometer (Sciex, Framingham, MA) was used coupled with a Dionex Ultimate 3000 UHPLC (ThermoFisher Scientific, Waltham, MA). Eight µl of sample was injected which represented 100 fmol of SIL labeled QconCAT peptides per 5 µg total protein per run. Data was analyzed with Skyline v4.2 software (MacCoss Lab Software, Seattle, WA), and was manually validated for transition quality, peak shape, and peak area boundaries. Savitsy-Golay smoothing was performed and the integrated peak areas were determined; data was further quantified by using a ratio of the $^{12}C$ peptide for the endogenous sample relative to the $^{13}C$ peptide from ECM targeted QconCATs. Linear range, limit of detection and limit of quantitation were obtained as previously defined (*55*). Previous studies were referenced to define the linear range, limit of detection, and limit of quantitation. Peptides were excluded from the analysis if they fell outside of the established parameters.

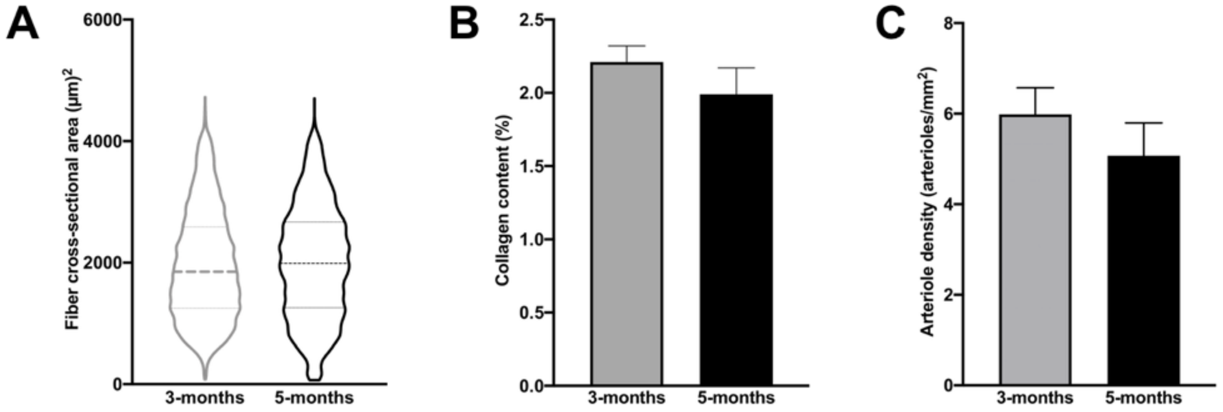


**Figure S1. Pelvic floor muscle fiber cross-sectional area, total collagen content, and overall arteriole density do not differ between 3- vs 5-month old rats.** (A) Violin plot for fiber cross-sectional area with median indicated by dash line. (B) Collagen content quantification. (C) Total vessel density. N=3/group. P-values derived from Student's t-tests for parametric and Mann-Whitney tests for non-parametric data, mean±SEM.

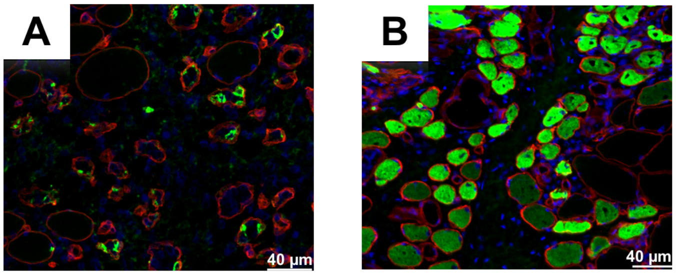


**Figure S2. Representative images of regenerating myofibers after simulated birth injury.** Fibers were identified by the presence of embryonic myosin heavy chain (green) at 3 (A) and 7 (B) days post-simulated birth injury.

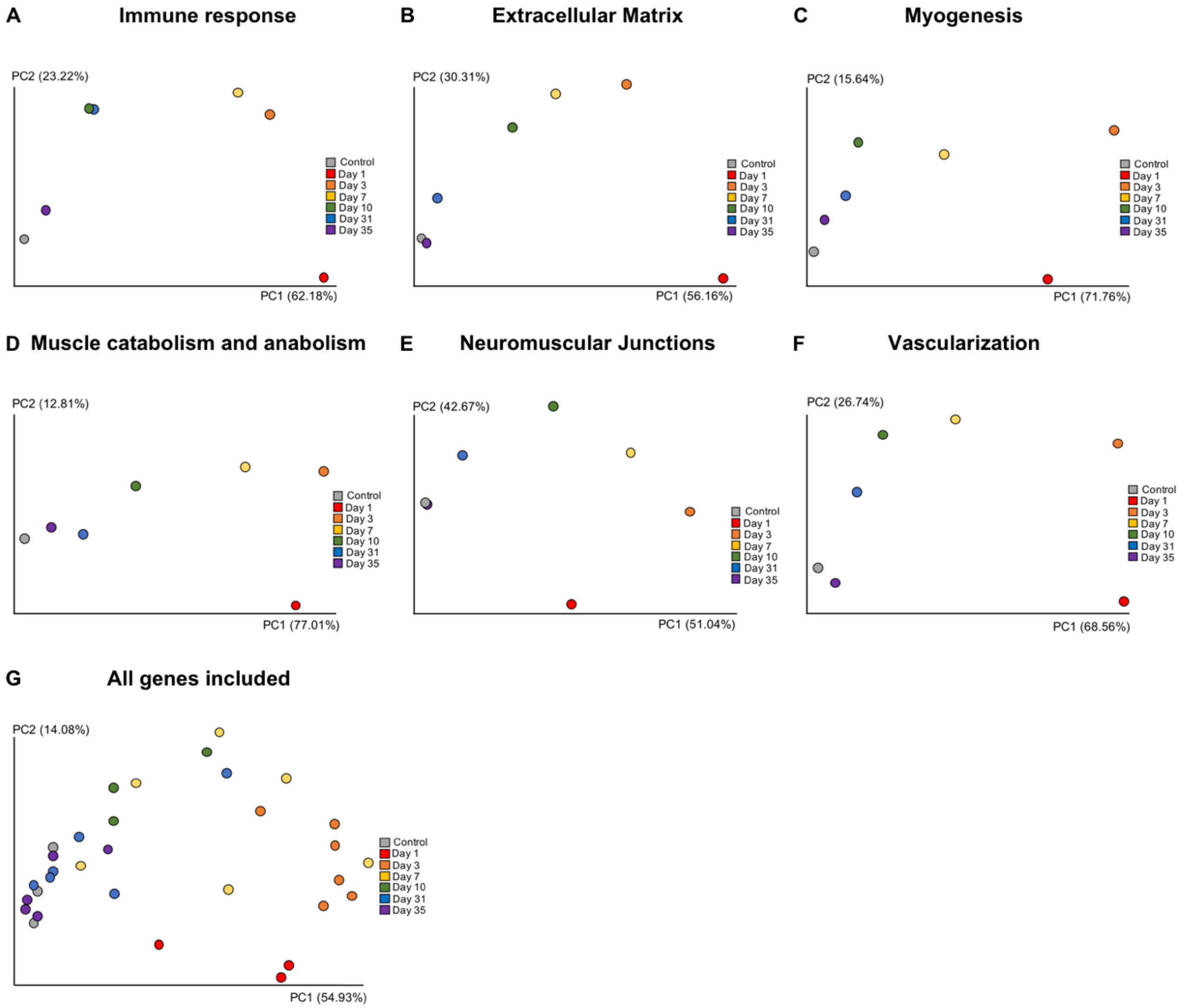


**Figure S3. Principal component analysis of transcriptional signatures during active regeneration of the pelvic floor muscles following simulated birth injury.** Customized Nanostring nCounter panel included genes involved in the following pathways: immune response (A), extracellular matrix (B), myogenesis (C), muscle anabolism and catabolism (D), neuromuscular junctions (E), vascularization (F). Average of normalized expression at each time point was included for all the pathways. (G) Principal component analysis for all genes including all replicates for each time point. Gene expression values were normalized to the following housekeeping genes Ap3d1, Hprt1, Rpl13a, Rpl32, Rplp0, and Tbp.

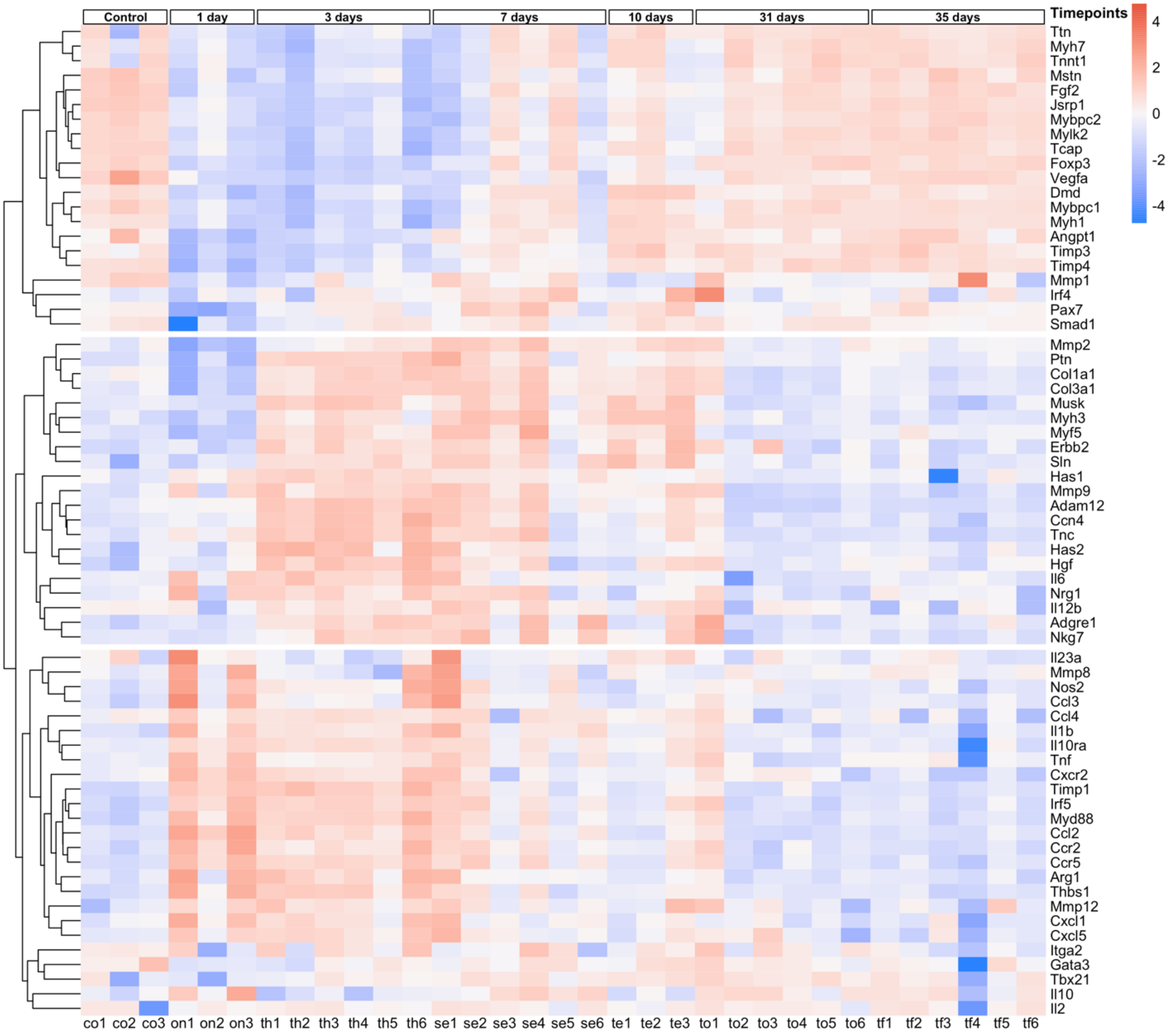


**Figure S4. Gene expression profile across the samples per timepoint post-simulated birth injury.** Heat map of genes with the highest variation in the dataset. N= 3/6 per group.

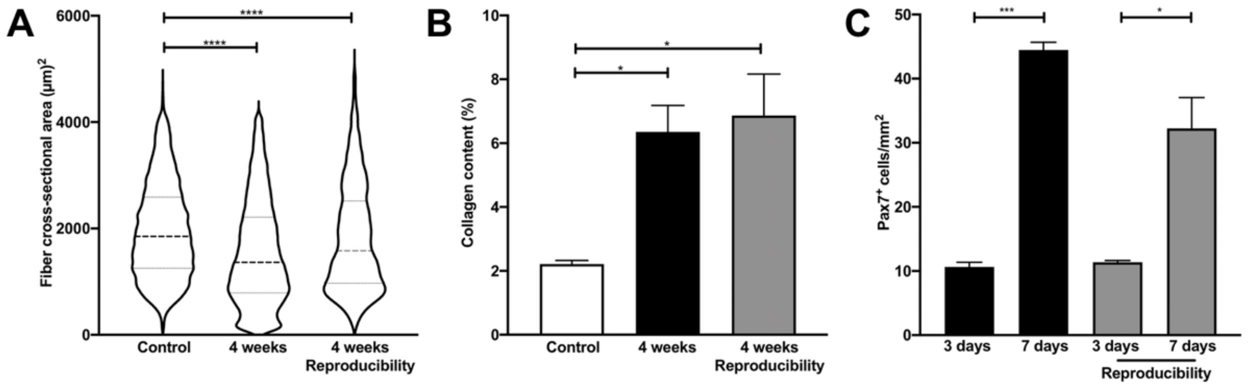


**Figure S5. Reproducibility of simulated birth injury (SBI).** A separate set of animals underwent SBI to assure the reproducibility of the main study outcomes. (A) Violin plot of fiber cross-sectional area. (B) Collagen content quantification. (C) Cell density of muscle stem cells. SBIs and quantification analyses for this set of animals were performed by the researchers not involved in the SBI experiments or analyses described in the main results. N=3-7/group. P-values derived from Student's t-tests or one-way ANOVA followed by pairwise comparisons with Tukey's range test for parametric and Mann-Whitney tests for non-parametric data.*p<0.05, ****p<0.0001, mean±SEM.

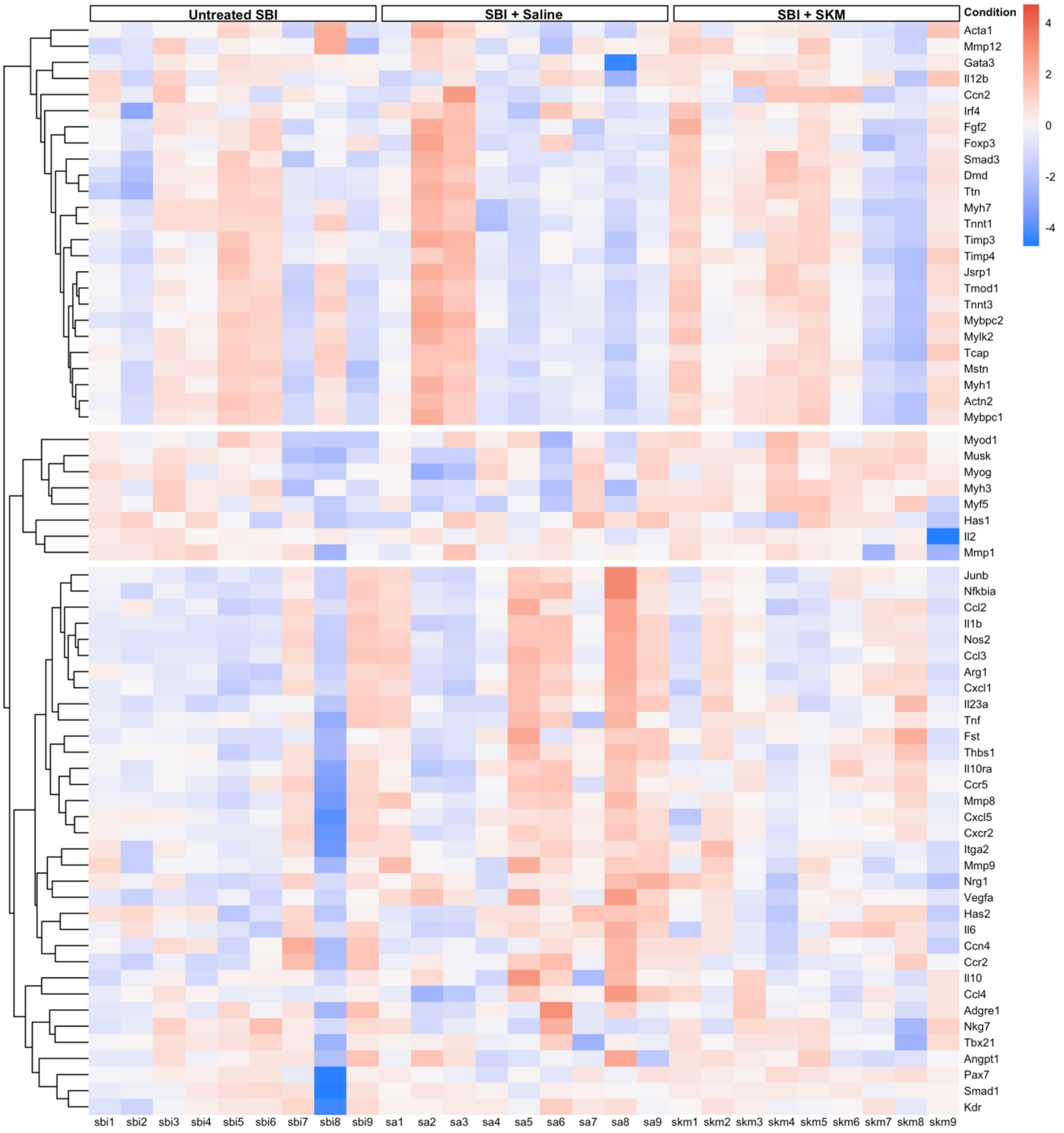


**Figure S6. Gene expression profile across the samples 3 days after immediate injection of skeletal muscle extracellular matrix (SKM) hydrogel.** Heat map of genes with the highest variation in the dataset. N=8/9 per group.

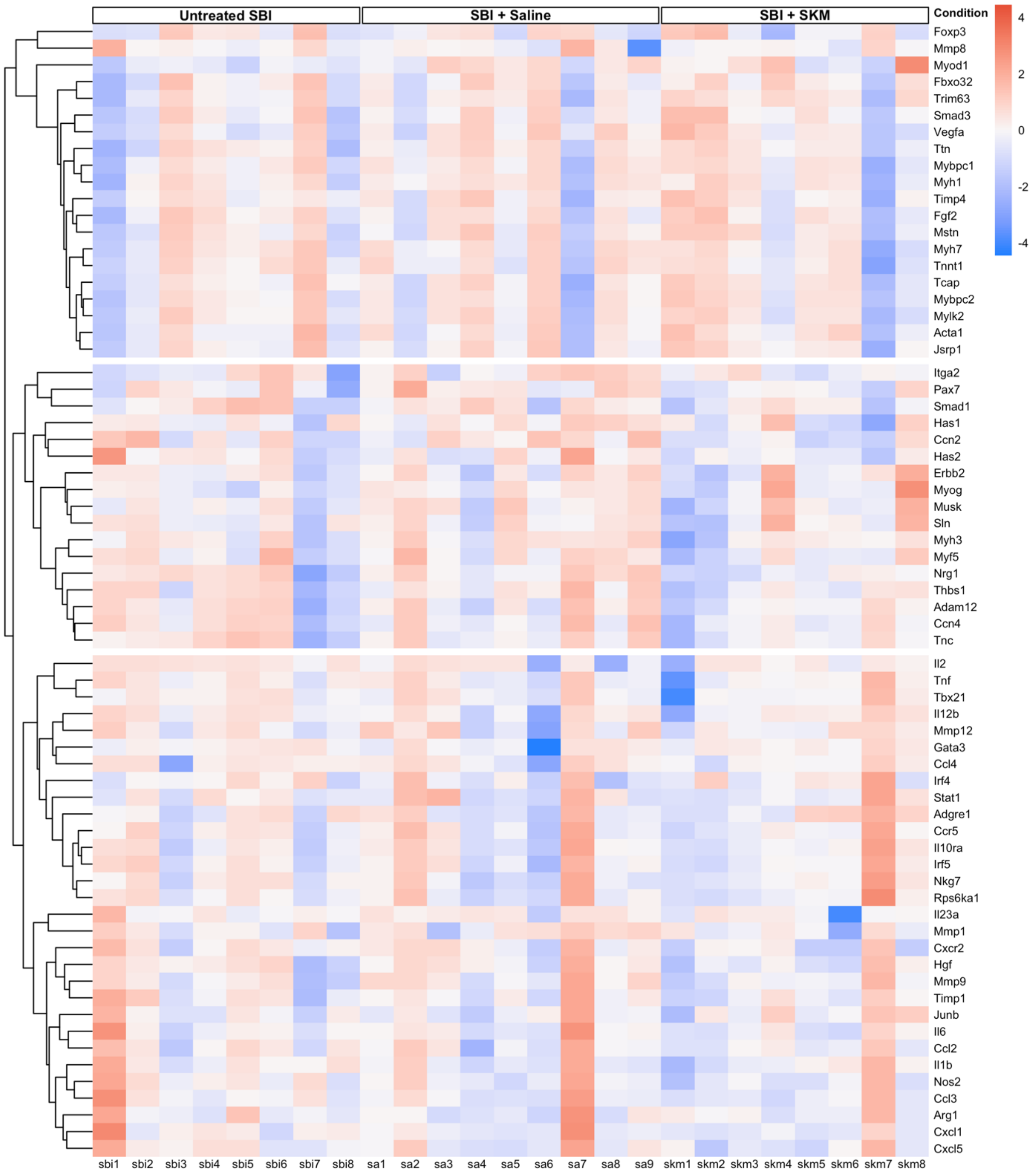


**Figure S7. Gene expression profile across the samples 7 days after immediate injection of skeletal muscle extracellular matrix (SKM) hydrogel.** Heat map of genes with the highest variation in the dataset. N=8/9 per group.

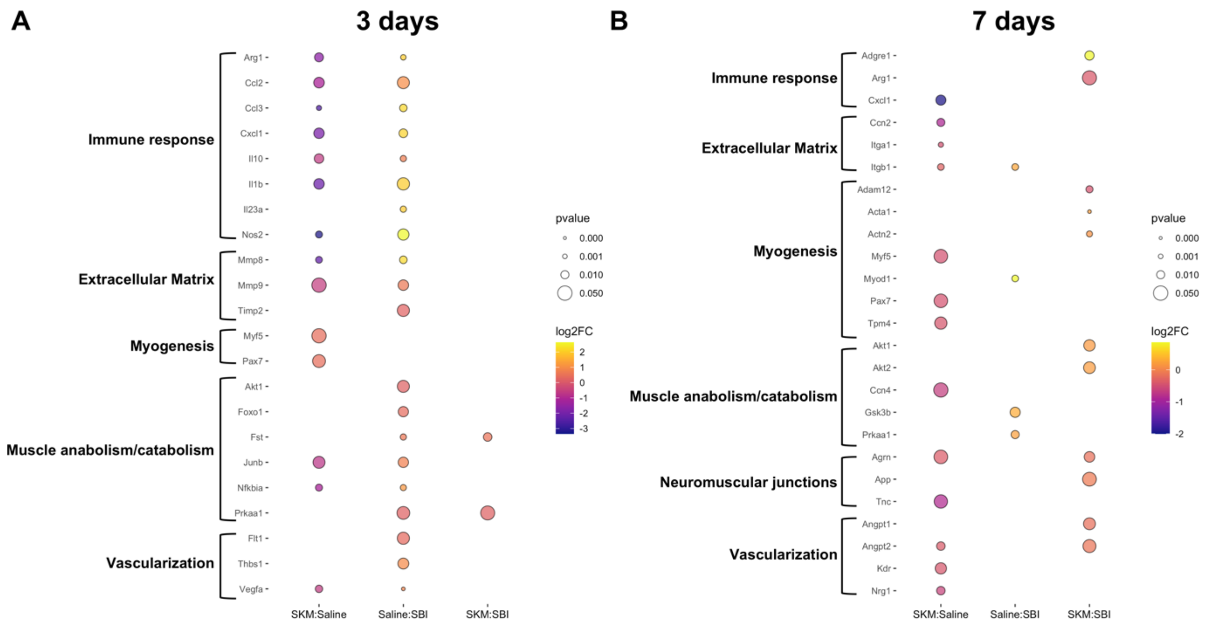


**Figure S8. Pairwise comparisons after immediate injection of skeletal muscle extracellular matrix (SKM) hydrogel.** Significantly differentially expressed genes at 3 (A) and 7 (B) days.

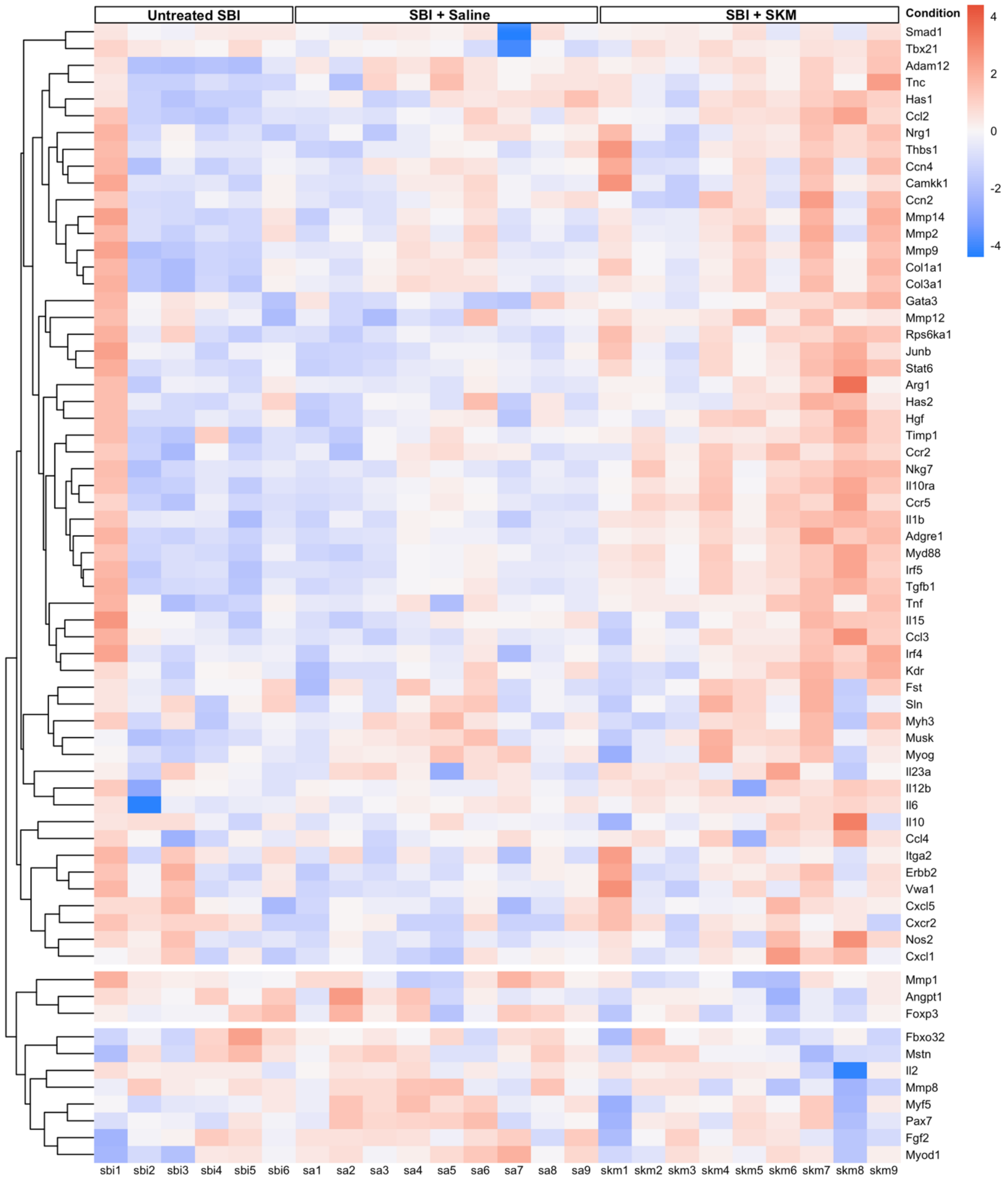


**Figure S9. Gene expression profile across the samples 3 days after delayed injection of skeletal muscle extracellular matrix (SKM) hydrogel.** Heat map of genes with the highest variation in the dataset. N=6/9 per group.

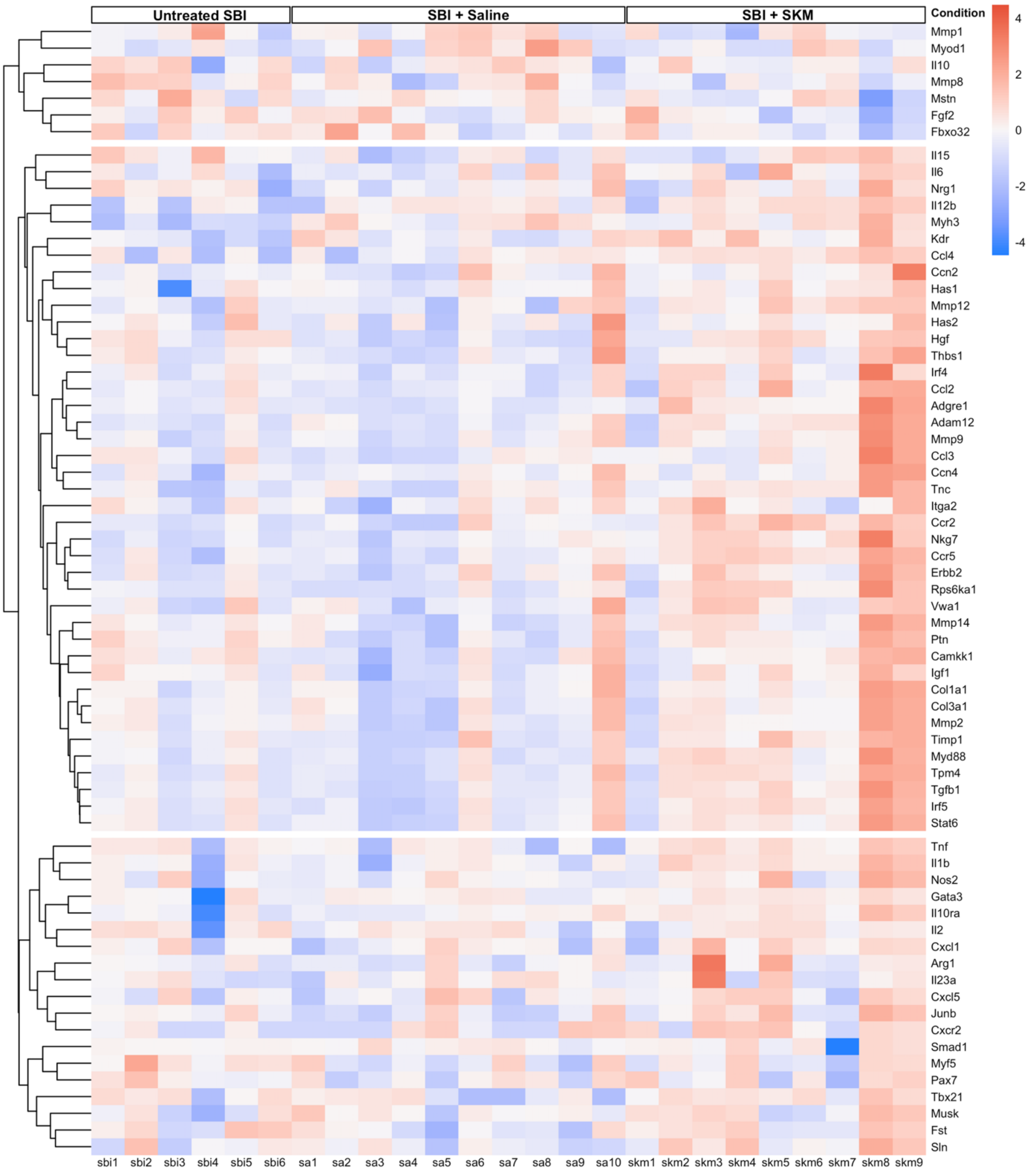


**Figure S10. Gene expression profile across the samples 7 days after delayed injection of skeletal muscle extracellular matrix (SKM) hydrogel.** Heat map of genes with the highest variation in the dataset. N=6/10 per group.

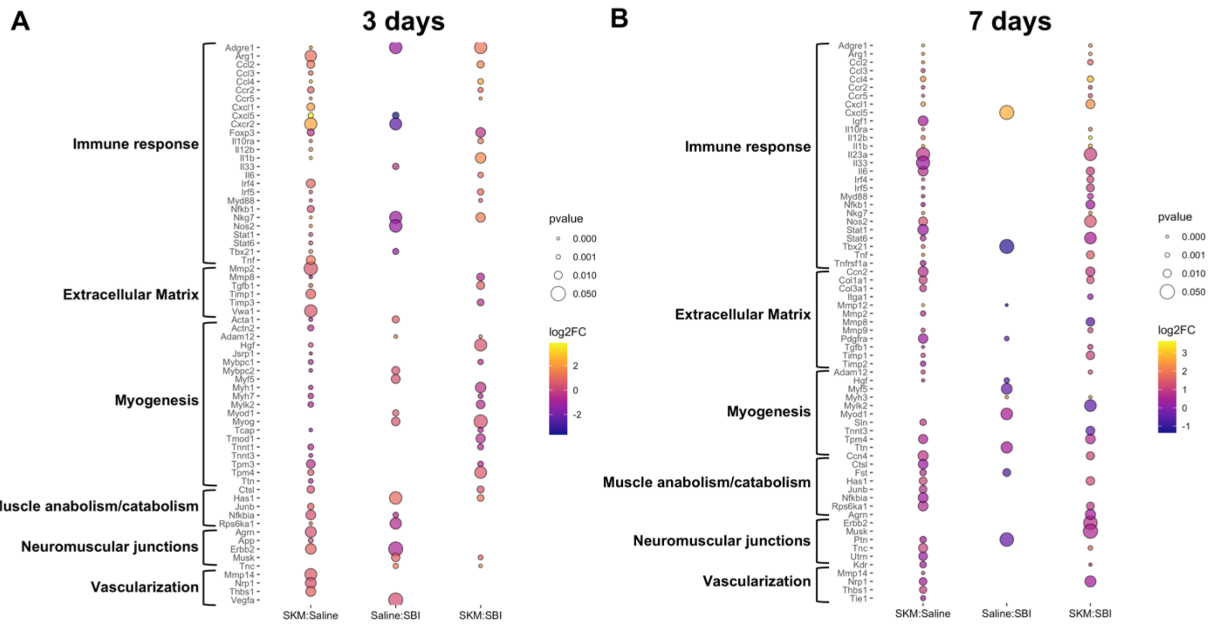


**Figure S11. Pairwise comparisons after delayed injection of skeletal muscle extracellular matrix (SKM) hydrogel.** Significantly differentially expressed genes at 3 (A) and 7 (B) days.

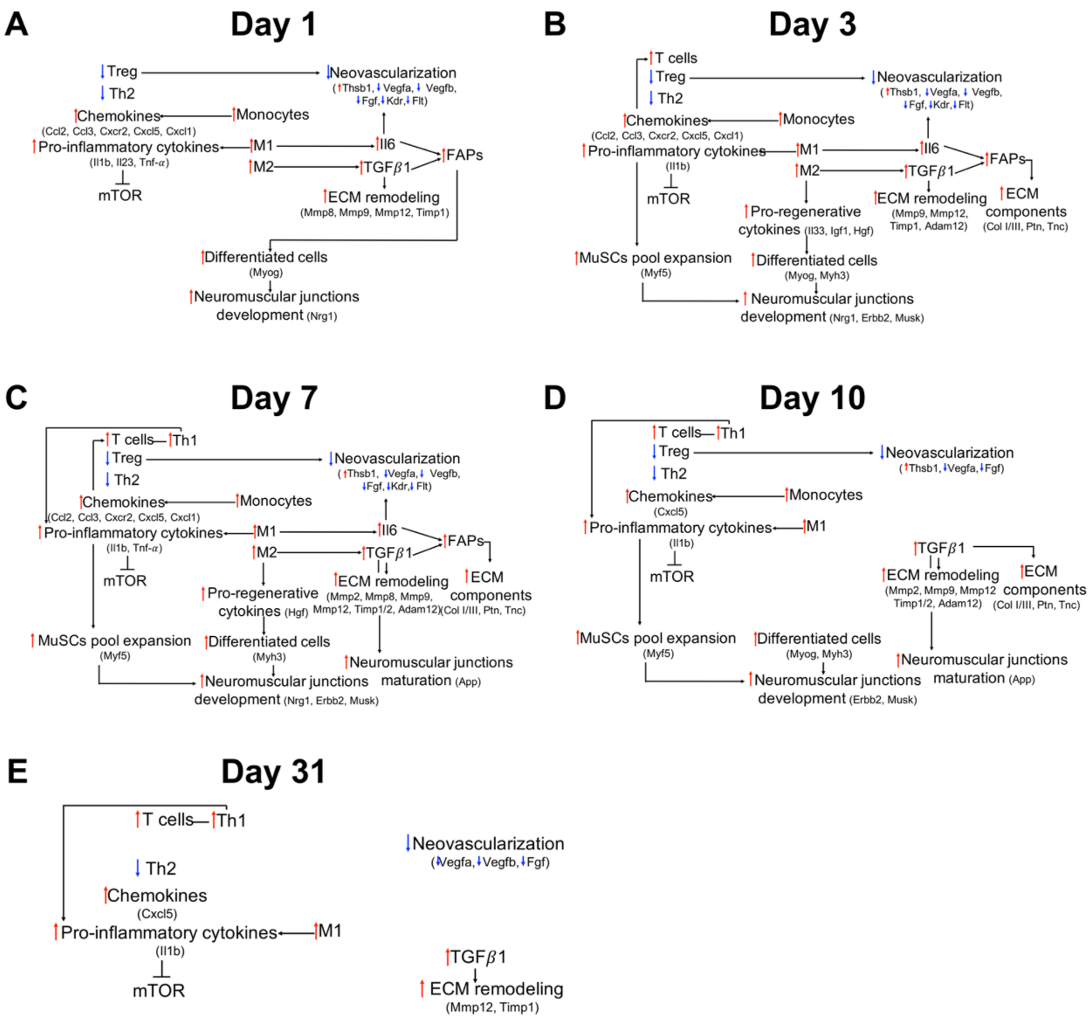


**Figure S12. Schematic representation of the interplay between different pathways analyzed at multiple time points following simulated birth injury.** 1 day (A), 3 days (B), 7 days (C), 10 days (D), 31 days (E).

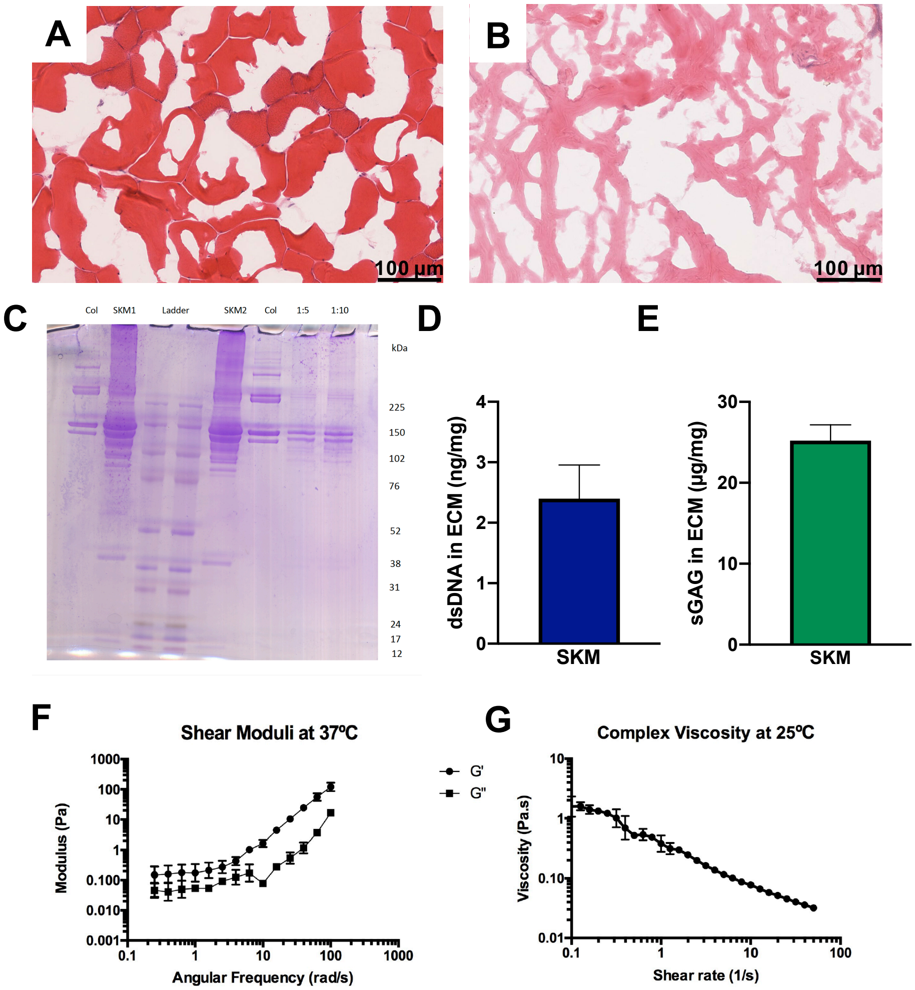


**Figure S13. Characterization of skeletal muscle extracellular matrix (SKM).** H&E of fresh porcine longissimus dorsi skeletal muscle (A), and decellularized SKM (B). C) SDS-PAGE of SKM replicates (SKM1, SKM2) and dilutions (1:5, 1:10) compared to a single ECM component, collagen (Col). The dsDNA (D) and the sGAG (E) composition in SKM (n=3). F, G) Mechanical properties of SKM, indicating storage (G') and loss (G") moduli for SKM (F) and complex viscosity traces of liquid SKM (G), indicating a shear thinning material.

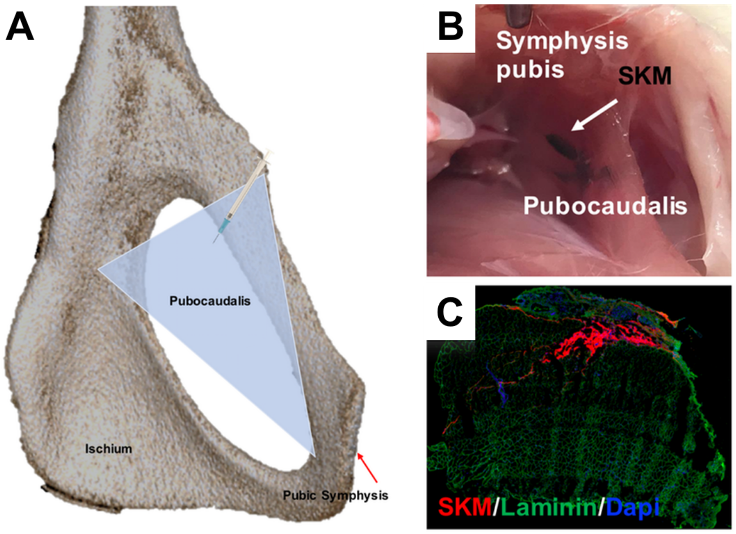


**Figure S14. Intramuscular injection of decellularized skeletal muscle extracellular matrix (SKM) hydrogel.** (A) Computer tomography image of the female rat pelvis demonstrating that SKM was delivered using the obturator foramen as a therapeutic window were the syringe passed through and injected the proximal (i.e., enthesial) region of pubocaudalis. We performed practice injections to assure a reliable delivery of SKM by prelabeling either with india ink (B) or Alexa Fluor 568 (C).

**Table S1. Customized Nanostring nCounter panel list.**

| Category | Gene | Accession | Classification |
|---|---|---|---|
| **Immune response Pro-inflammatory** | Il1b | NM_031512.1 | Cytokine |
| | Il6 | NM_053836.1 | Cytokine |
| | Tnf | NM_012675.2 | Cytokine |
| | Ifng* | NM_138880.2 | Cytokine |
| | Il12b | NM_022611.1 | Cytokine |
| | Il2 | NM_022611.1 | Cytokine |
| | Tnfrsf1a | NM_013091.1 | Receptor/growth factor |
| | Ccl2 | NM_031530.1 | Chemokine |
| | Ccl3 | NM_013025.2 | Chemokine |
| | Ccl4 | NM_053858.1 | Chemokine |
| | Ccr2 | NM_021866.1 | Chemokine |
| | Cxcl1 | NM_053960.3 | Chemokine |
| | Cxcl5 | NM_053304.1 | Chemokine |
| | Cxcr2 | NM_017183.1 | Chemokine |
| | Nos2 | NM_012611.2 | M1 |
| | Tbx21 | NM_001271120.1 | Th1 |
| | Il23a | NM_130410.2 | Cytokine |
| | Stat1 | NM_032612.2 | M1 transcription factor |
| | Irf5 | NM_001106586.1 | M1 transcription factor |
| | Adgre1 | NM_001007557.1 | Macrophages |
| | Myd88 | NM_198130.1 | Monocytes |
| | Nfkb1 | XM_342346.3 | Transcription factor |
| | Nkg7 | NM_133540.1 | T cells |
| | Mmp12 | NM_053963.2 | Metalloelastase |
| **Immune response Pro-regenerative (Anti-inflammatory)** | Il33 | NM_001014166.1 | Cytokine |
| | Il13* | NM_053828.1 | Cytokine |
| | Il4* | NM_201270.1 | Cytokine |
| | Il5* | NM_021834.1 | Cytokine |
| | Il15* | XM_008772363.2 | Cytokine |
| | Il17* | NM_001106897.1 | Cytokine |
| | Il10 | NM_012854.2 | Cytokine |

| | | | |
|---|---|---|---|
| | Il10ra | NM_057193.2 | Cytokine/Receptor |
| | Igf1 | NM_001082477.2 | Growth factor |
| | Igf1r | NM_052807.2 | Growth factor/Receptor |
| | Ccr5 | NM_053960.3 | Chemokine |
| | Arg1 | NM_017134.2 | M2 |
| | Gata3 | NM_133293.1 | Th2 |
| | Foxp3 | NM_001108250.1 | T regulatory cells |
| | Stat6 | NM_001044250.1 | M2 transcription factor |
| | Irf4 | NM_001106108.1 | M2 transcription factor |
| **Myogenesis** | Pax7 | NM_001191984.1 | Quiescent and activated muscle stem cells |
| | Myod1 | NM_176079.1 | Activated muscle stem cells |
| | Myf5 | NM_001106783.1 | Activated cells |
| | Myf6 | NM_013172.1 | Differentiated cells |
| | Myog | NM_017115.2 | Differentiated muscle stem cells |
| | MyH3 | NM_012604.1 | Regenerated myofibers |
| | Hgf | NM_017017.2 | Growth factor |
| | Adam12 | XM_017590240.1 | Metallopeptidase |
| | Actn2 | NM_001170325.1 | Muscle structure |
| | Des | NM_022531.1 | Muscle structure |
| | Dmd | NM_001005244.1 | Muscle structure |
| | Tpm3 | NM_173111.1 | Muscle structure |
| | Tpm4 | NM_012678.2 | Muscle structure |
| | Tcap | NM_001271277.1 | Muscle structure |
| | Acta1 | NM_019212.2 | Muscle structure |
| | Mybpc1 | NM_001100758.1 | Muscle structure |
| | Mybpc2 | NM_001106257.1 | Muscle structure |
| | Tmod1 | NM_013044.2 | Muscle structure |
| | Ttn | XM_001065955.4 | Muscle structure |
| | Jsrp1 | NM_001109591.1 | Muscle structure |
| | Tnnt1 | NM_001277260.1 | Muscle structure |
| | Tnnt3 | NM_001270665.1 | Muscle structure |
| | MyH1 | NM_001135158.1 | Muscle structure |
| | MyH7 | NM_017240.1 | Muscle structure |
| | Mylk2 | NM_057209.1 | Muscle structure |
| | Sln | NM_001013247.1 | Muscle structure |

| | | | |
|---|---|---|---|
| | Wnt7a* | NM_001100473.1 | Muscle homeostasis |
| Extracellular Matrix | Mmp1 | NM_001134530.1 | Metalloproteinase |
| | Mmp8 | NM_053963.2 | Metalloproteinase |
| | Mmp13* | NM_133530.1 | Metalloproteinase |
| | Mmp2 | NM_031054.2 | Metalloproteinase |
| | Mmp9 | NM_031055.1 | Metalloproteinase |
| | Timp1 | NM_053819.1 | Tissue inhibitor of metalloproteinase |
| | Timp2 | NM_021989.2 | Tissue inhibitor of metalloproteinase |
| | Timp3 | NM_012886.2 | Tissue inhibitor of metalloproteinase |
| | Timp4 | NM_001109393.1 | Tissue inhibitor of metalloproteinase |
| | Ccn2 | NM_022266.2 | Growth factor |
| | Pdgfa | NM_012801.1 | Growth factor |
| | Pdgfra | NM_012802.1 | Growth factor/Receptor |
| | Dcn | NM_024129.1 | Proteoglycan-ECM component |
| | Tgfb1 | NM_021578.2 | Growth factor |
| | Tgfbr1 | NM_012775.2 | Growth factor/Receptor |
| | Smad2 | NM_001105817.1 | Growth factor/Receptor |
| | Smad3 | NM_013095.2 | Signal transducer |
| | Smad4 | NM_019275.2 | Signal transducer |
| | Col1a1 | NM_053304.1 | ECM component |
| | Col3a1 | NM_032085.1 | ECM component |
| | Itga1 | NM_030994.2 | ECM component |
| | Itgb1 | NM_017022.2 | Cell-matrix interaction |
| | Itga2 | XM_345156.6 | Cell-matrix interaction |
| | Vwa1 | NM_001013938.1 | Cell-matrix interaction |
| | Akt1 | NM_033230.1 | Anabolism/mTOR signaling |
| | Mtor | NM_019906.1 | Anabolism/mTOR signaling |
| | Rheb | NM_013216.1 | Anabolism/mTOR signaling |
| | Eif2b5 | NM_138866.2 | Anabolism/mTOR signaling |
| | Akt2 | NM_017093.1 | Anabolism/mTOR signaling |
| | Eif4b | NM_001008324.1 | Anabolism/mTOR signaling |
| | Eif4e | NM_053974.2 | Anabolism/mTOR signaling |
| | Eif4ebp1 | NM_053857.1 | Anabolism/mTOR signaling |
| | Eif4g1 | XM_213569.6 | Anabolism/mTOR signaling |

| | | | |
|---|---|---|---|
| **Muscle anabolism and catabolism** | Grb10 | NM_001109093.1 | Anabolism |
| | Rps6ka1 | NM_031107.1 | Anabolism |
| | Rps6kb1 | NM_031985.1 | mTOR signaling |
| | Gsk3b | NM_032080.1 | Anabolism |
| | Junb | NM_021836.2 | Anabolism |
| | Nol3 | NM_053516.2 | Apoptosis |
| | Ikbkb | NM_053355.1 | Anabolism |
| | Nfkbia | XM_001075778.1 | Anabolism |
| | Foxo1 | NM_001191846.2 | Catabolism |
| | Foxo3 | NM_001106395.1 | Catabolism |
| | Fbxo32 | NM_133521.1 | Catabolism |
| | Trim63 | NM_080903.1 | Catabolism |
| | Camkk1 | NM_031662.1 | Catabolism |
| | Mstn | NM_019151.1 | Anabolism |
| | Ctsl | NM_013156.2 | Catabolism |
| | Prkaa1 | NM_019142.2 | Catabolism |
| | Fst | NM_012561.1 | Anabolism |
| | Ccn4 | NM_031716.1 | Cellular survival |
| | Has1 | NM_172323.1 | Cellular survival |
| | Has2 | NM_013153.1 | Cellular survival |
| **Angiogenesis** | Vegfa | NM_031836.2 | Growth factor |
| | Vegfb | NM_053549.1 | Growth factor |
| | Angpt1 | NM_053546.1 | Growth factor |
| | Angpt2 | NM_134454.1 | Growth factor |
| | Kdr | NM_013062.1 | Growth factor/Receptor |
| | Flt1 | NM_019306.1 | Growth factor/Receptor |
| | Nrp1 | NM_145098.2 | Growth factor/Receptor |
| | Tie1 | NM_053545.1 | Growth factor/Receptor |
| | Fgf2 | NM_019305.2 | Growth factor/Receptor |
| | Thsb1 | NM_001013062.1 | Cell-matrix interaction |
| | Mmp14 | NM_031056.1 | Metalloproteinase |
| | Erbb2 | NM_017003.2 | Growth factor/Receptor |
| | Nrg1 | NM_001271120.1 | Growth factor |
| | App | NM_019288.1 | Receptor |
| | Dvl1 | NM_031820.1 | Cytoplasmic |

| | | | |
|---|---|---|---|
| **Neuromuscular Junctions development and maintenance** | Ptn | NM_017066.2 | Growth factor |
| | Tnc | NM_053861.1 | ECM component |
| | Utrn | NM_013070.1 | Cytoskeleton |
| | Musk | NM_031061.1 | Receptor |
| | Agrn | NM_175754.1 | Proteoglycan |
| | Gphn | NM_022865.3 | Cytoskeleton |
| **Housekeeping genes** | Ap3d1 | XM_001076208.1 | |
| | Hprt1 | NM_012583.2 | |
| | Rpl13a | NM_173340.2 | |
| | Rpl32 | NM_013226.2 | |
| | Rplp0 | NM_022402.2 | |
| | Tbp | NM_001004198.1 | |

Note: *Genes that were excluded since normalized mRNA counts were at the level of background for all conditions.

**Table S2. Immune response related genes throughout the different time points after simulated birth injury**. P and q values are indicated for significantly differentially expressed genes compared to uninjured group. FC= Log2FC

| Days after Simulated Birth Injury | | | | | | | | | | | | | | | | | | |
|---|---|---|---|---|---|---|---|---|---|---|---|---|---|---|---|---|---|---|
| **Gene** | 1 | | | 3 | | | 7 | | | 10 | | | 31 | | | 35 | | |
| | FC | p | q | FC | p | q | FC | p | q | FC | p | q | FC | p | q | FC | p | q |
| **Myd88** | 2.13 | 2.60E-08 | 1.80E-07 | 1.94 | 2.80E-08 | 1.20E-07 | 1.23 | 0.0002 | 0.0009 | 0.77 | 0.03 | 0.1 | 0.58 | | | 0.26 | | |
| **Adgre1** | 0.38 | | | 2.6 | 0.001 | 0.002 | 2.85 | 0.0005 | 0.001 | 1.68 | 0.04 | 0.12 | 2.27 | 0.004 | 0.07 | 0.93 | | |
| **Nkg7** | -0.26 | | | 1.95 | 0.01 | 0.01 | 2.57 | 0.001 | 0.003 | 1.75 | 0.03 | 0.1 | 1.63 | 0.02 | 0.1 | -0.35 | | |
| **Foxp3** | -3.07 | 8.40E-01 | 9.70E-09 | -3.66 | 0 | 0 | -0.99 | 0.009 | 0.01 | -1.27 | 0.006 | 0.04 | -0.17 | | | -0.18 | | |
| **Nos2** | 6.99 | 5.90E-05 | 0.0001 | 5.06 | 2.20E-08 | 9.70E-08 | 5.81 | 1.40E-11 | 1.90E-09 | 0.75 | | | 1.8 | | | 0.44 | | |
| **Stat1** | 0.25 | | | 0.44 | | | 0.78 | 0.001 | 0.003 | 0.6 | 0.02 | 0.1 | 0.44 | | | 0.28 | | |
| **Irf5** | 2.67 | 5.70E-07 | 2.60E-06 | 2.45 | 7.30E-07 | 1.90E-06 | 1.92 | 6.20E-05 | 0.0003 | 1.34 | 0.008 | 0.055 | 1.1 | 0.01 | 0.1 | 0.59 | | |
| **Tbx21** | -1.83 | 0.01 | 0.02 | 0.48 | | | 1.23 | 0.03 | 0.055 | 1.47 | 0.04 | 0.1 | 1.65 | 0.006 | 0.08 | 0.77 | | |
| **Il1b** | 7.52 | 2.10E-06 | 7.80E-06 | 5.22 | 0.0001 | 0.0002 | 6.6 | 9.00E-06 | 0.0001 | 2.64 | 0.03 | 0.11 | 2.72 | 0.02 | 0.1 | 1.06 | | |
| **Il23** | 1.82 | 0.03 | 0.04 | -0.72 | | | 0.84 | | | 0.54 | | | -0.48 | | | -0.48 | | |
| **Il2** | -0.06 | | | 0.23 | | | 0.42 | | | -0.14 | | | 0.24 | | | 0.34 | | |
| **Il15** | -1.4 | 9.20E-07 | 4.00E-06 | -0.32 | | | 0.06 | | | 0.03 | | | -0.12 | | | 0.23 | | |
| **Il12b** | -0.59 | | | 1.15 | | | 2.21 | 0.007 | 0.01 | 1.34 | | | 1.02 | | | -1.29 | | |
| **Il6** | 3.96 | 0.0001 | 0.0003 | 4.11 | 5.40E-05 | 0.001 | 2.72 | 0.003 | 0.007 | 0.24 | | | -0.94 | | | 0.01 | | |
| **Tnf** | 3.61 | 2.00E-05 | 6.20E-05 | 1.4 | | | 1.89 | 0.009 | 0.01 | 0.99 | | | 1.11 | | | 0.59 | | |
| **Tnfrsf1a** | 1.02 | 2.80E-05 | 8.20E-05 | 1.01 | 4.00E-06 | 9.80E-06 | 0.37 | | | 0.24 | | | 0.13 | | | -0.04 | | |
| **Cxcl5** | 5.83 | 0.001 | 0.003 | 5.51 | 0.001 | 0.003 | 6.07 | 9.40E-11 | 3.20E-09 | 2.83 | | | 3.82 | 0.01 | 0.1 | 1.15 | | |
| **Ccl4** | 2.6 | 0.001 | 0.002 | 1.61 | 0.01 | 0.02 | 1.73 | 0.01 | 0.02 | 0.63 | | | 0.38 | | | -0.88 | | |
| **Ccr2** | 2.99 | 5.50E-08 | 3.10E-07 | 1.94 | 7.10E-05 | 0.0001 | 1.54 | 0.001 | 0.003 | 0.99 | | | 0.82 | | | 0.026 | | |
| **Cxcr2** | 6.9 | 1.30E-05 | 4.30E-05 | 4.32 | 0.001 | 0.002 | 3.71 | 7.20E-11 | 3.20E-09 | -0.17 | | | 1.82 | | | -1.88 | | |
| **Ccl2** | 6.98 | 1.60E-11 | 4.50E-10 | 4.14 | 2.40E-06 | 6.20E-06 | 2.83 | 0.0004 | 0.001 | 1.12 | | | 0.98 | | | 0.31 | | |
| **Ccl3** | 5.29 | 1.10E-06 | 4.60E-06 | 2.6 | 0.004 | 0.006 | 3.61 | 0.0002 | 0.0009 | 0.89 | | | 1.24 | | | 0.81 | | |
| **Cxcl1** | 5.68 | 0.0001 | 0.003 | 3.51 | 0.006 | 0.008 | 3.43 | 0.007 | 0.01 | 0.89 | | | 0.06 | | | 0.14 | | |
| **Mmp12** | 5.59 | 0.01 | 0.01 | 6.17 | 0.006 | 0.008 | 5.5 | 0.008 | 0.01 | 7.59 | 1.40E-08 | 1.00E-06 | 5.38 | 0.009 | 0.1 | 5.08 | 3.50E-12 | 4.90E-10 |
| **Nfkb1** | 0.11 | | | 0.29 | | | 0.14 | | | 0.09 | | | 0.17 | | | -0.01 | | |
| **Arg1** | 4.04 | 2.60E-06 | 9.40E-06 | 2.98 | 0.0001 | 0.0002 | 2 | 0.006 | 0.1 | 0.4 | | | 0.26 | | | -0.09 | | |
| **Gata3** | -2.68 | 1.10E-05 | 3.80E-05 | -1.58 | 0.0003 | 0.0006 | -1.04 | 0.01 | 0.02 | -0.67 | | | -0.93 | 0.04 | 0.1 | -1.32 | 0.01 | 0.25 |
| **Stat6** | 1.33 | 4.40E-05 | 0.0001 | 1.27 | 1.70E-05 | 3.90E-05 | 0.85 | 0.003 | 0.006 | 0.55 | | | 0.59 | 0.03 | 0.1 | 0.26 | | |
| **Irf4** | -0.16 | | | 0.24 | | | 0.74 | | | 0.82 | | | 0.84 | | | 0.12 | | |
| **Igf1** | -1.06 | 0.01 | 0.02 | 1.04 | 0.007 | 0.01 | 0.64 | | | 0.31 | | | -0.08 | | | -0.19 | | |
| **Igf1r** | 0.46 | | | 0.47 | 0.03 | 0.04 | 0.42 | | | 0.72 | 0.003 | 0.03 | 0.55 | 0.01 | 0.1 | 0.24 | | |
| **Il10** | 1.1 | 0.0003 | 0.0006 | -0.19 | | | 0.68 | 0.01 | 0.02 | -0.05 | | | 0.63 | 0.02 | 0.1 | 0.39 | | |
| **Il10ra** | 2.96 | 4.70E-05 | 0.0001 | 2.71 | 5.20E-05 | 0.0001 | 2.04 | 0.001 | 0.003 | 1.52 | 0.02 | 0.1 | 1.17 | | | -0.07 | | |
| **Il33** | -0.44 | | | 0.72 | 0.001 | 0.001 | 0.25 | | | 0.33 | | | 0.7 | 0.001 | 0.03 | 0.49 | 0.02 | 0.44 |
| **Ccr5** | 3.84 | 1.10E-07 | 6.10E-07 | 3.05 | 4.20E-06 | 1.00E-05 | 2.49 | 0.0001 | 0.0005 | 1.38 | 0.03 | 0.1 | 1.16 | | | 0.15 | | |

**Table S3. Myogenesis related genes throughout the different time points after simulated birth injury.** P and q values are indicated for significantly differentially expressed genes compared to uninjured group. Log2FC

| | Days after Simulated Birth Injury | | | | | | | | | | | | | | | | | |
|---|---|---|---|---|---|---|---|---|---|---|---|---|---|---|---|---|---|---|
| **Gene** | 1 | | | 3 | | | 7 | | | 10 | | | 31 | | | 35 | | |
| | FC | p | q | FC | p | q | FC | p | q | FC | p | q | FC | p | q | FC | p | q |
| **Pax7** | -2.19 | 2.10E-08 | 1.50E-07 | -0.08 | | | 0.46 | | | 0.58 | | | -0.16 | | | 0.12 | | |
| **Myf5** | -0.89 | 0.02 | 0.03 | 1.12 | 0.001 | 0.002 | 1.48 | 2.70E-05 | 0.0002 | 1.34 | 0.0005 | 0.01 | 0.05 | | | 0.49 | | |
| **Myod1** | -2.13 | 3.00E-08 | 2.00E-07 | -1.62 | 1.40E-07 | 4.90E-07 | -1.8 | 4.70E-09 | 9.60E-08 | -0.39 | | | -1.26 | 4.60E-05 | 0.003 | -1.09 | 0.0004 | 0.01 |
| **Myf6** | 0.25 | | | -1.5 | 8.20E-09 | 4.30E-08 | -0.64 | 0.01 | 0.02 | 0.08 | | | -0.75 | 0.004 | 0.07 | -0.58 | 0.02 | 0.4 |
| **Myog** | 1.09 | 0.001 | 0.003 | 1.65 | 3.20E-07 | 9.80E-07 | 0.32 | | | 1.01 | 0.003 | 0.03 | -0.32 | | | -0.14 | | |
| **Myh3** | -1.96 | 0.03 | 0.04 | 3.51 | 0.0001 | 0.0002 | 5.6 | 3.60E-08 | 6.20E-07 | 6.11 | 1.40E-08 | 1.00E-06 | 1.38 | | | -0.27 | | |
| **Hgf** | 0.43 | | | 1.73 | 9.30E-07 | 2.40E-06 | 0.94 | 0.006 | 0.01 | 0.96 | 0.01 | 0.07 | 0.65 | | | 0.53 | | |
| **Adam12** | 0.87 | | | 3.78 | 1.60E-07 | 5.50E-07 | 2.88 | 2.50E-05 | 0.0002 | 1.64 | 0.01 | 0.1 | -0.46 | | | -1.08 | | |
| **Acta1** | -1.37 | 0.02 | 0.03 | -2.94 | 6.50E-09 | 3.60E-08 | -1.3 | 0.01 | 0.01 | -1.14 | | | -0.77 | | | -0.21 | | |
| **Actn2** | -1.69 | 0.0002 | 0.0005 | -1.74 | 3.30E-06 | 8.20E-06 | -0.4 | | | 0.57 | | | -0.07 | | | -0.24 | | |
| **Des** | -0.76 | 0.006 | 0.01 | -1.01 | 1.70E-05 | 3.90E-05 | -0.46 | | | -0.16 | | | -0.27 | | | -0.08 | | |
| **Dmd** | -2.15 | 1.80E-06 | 6.80E-06 | -1.7 | 2.10E-06 | 5.50E-06 | -0.4 | | | 0.72 | | | -0.09 | | | -0.07 | | |
| **Jsrp1** | -1.67 | 0.0001 | 0.0004 | -2.51 | 6.60E-12 | 7.60E-11 | -1.33 | 0.0002 | 0.0009 | -0.96 | 0.02 | 0.1 | -0.66 | | | -0.33 | | |
| **Mybpc1** | -2.25 | 2.70E-05 | 8.20E-05 | -3.06 | 1.10E-12 | 1.60E-11 | -1.1 | 0.01 | 0.01 | -0.24 | | | -0.41 | | | -0.35 | | |
| **Mybpc2** | -2 | 0.002 | 0.003 | -2.87 | 8.70E-08 | 3.30E-07 | -1.26 | 0.01 | 0.02 | -0.81 | | | -0.63 | | | -0.14 | | |
| **Myh1** | -2.24 | 0.0003 | 0.0006 | -2.84 | 1.40E-08 | 7.00E-08 | -0.6 | | | 0.48 | | | 0.11 | | | 0.1 | | |
| **Myh7** | -1.62 | 0.02 | 0.03 | -2.4 | 6.10E-05 | 0.0001 | -0.28 | | | 0.4 | | | 0.6 | | | 0.56 | | |
| **Mylk2** | -2.64 | 0.0003 | 0.0006 | -3.42 | 5.10E-09 | 2.90E-08 | -1.28 | 0.02 | 0.03 | -1.15 | | | -0.32 | | | -0.01 | | |
| **Sln** | -0.04 | | | 2.46 | 3.00E-05 | 6.50E-05 | 2.76 | 4.40E-06 | 5.60E-05 | 3.65 | 4.40E-08 | 2.00E-06 | 0.88 | | | 0.43 | | |
| **Tcap** | -1.96 | 0.0005 | 0.0009 | -2.83 | 7.00E-10 | 5.70E-09 | -1.35 | 0.003 | 0.006 | -0.52 | | | -0.4 | | | -0.19 | | |
| **Tmod1** | -2.27 | 6.10E-07 | 2.70E-06 | -2.5 | 5.40E-12 | 6.80E-11 | -1.09 | 0.002 | 0.005 | -0.59 | | | -0.41 | | | -0.21 | | |
| **Tnnt1** | -1.11 | | | -1.91 | 5.80E-05 | 0.0001 | -0.31 | | | 0.24 | | | 0.47 | | | 0.5 | | |
| **Tnnt3** | -1.97 | 0.0001 | 0.0003 | -3.43 | 5.50E-16 | 1.50E-14 | -1.29 | 0.001 | 0.004 | -0.62 | | | -0.51 | | | -0.2 | | |
| **Tpm3** | -0.75 | | | -1.73 | 9.50E-05 | 0.0001 | -0.5 | | | 0.03 | | | 0.53 | | | 0.53 | | |
| **Tpm4** | 1.4 | 0.0002 | 0.0004 | 1.98 | 3.40E-08 | 1.40E-07 | 1.09 | 0.001 | 0.003 | 0.85 | 0.02 | 0.1 | 0.57 | | | 0.15 | | |
| **Ttn** | -2.69 | 0.006 | 0.009 | -2.63 | 0.0006 | 0.001 | -0.71 | | | 0.2 | | | 0.06 | | | | | |

**Table S4. Muscle anabolism and catabolism related genes throughout the different time points after simulated birth injury.** P and q values are indicated for significantly differentially expressed genes compared to uninjured group. Log2FC

| | **Days after Simulated Birth Injury** | | | | | | | | | | | | | | | | | |
|---|---|---|---|---|---|---|---|---|---|---|---|---|---|---|---|---|---|---|
| **Gene** | 1 | | | 3 | | | 7 | | | 10 | | | 31 | | | 35 | | |
| | FC | p | q | FC | p | q | FC | p | q | FC | p | q | FC | p | q | FC | p | q |
| **Has1** | 1.73 | | | 2.76 | 1.90E-07 | 6.60E-07 | 1.72 | 0.001 | 0.002 | 0.89 | | | 0.41 | | | 0.33 | | |
| **Ccn4** | 0.59 | | | 3.1 | 6.20E-08 | 2.40E-07 | 2.01 | | | 1.46 | 0.01 | 0.06 | 0.44 | | | -0.19 | | |
| **Has2** | 0.57 | | | 2.78 | 6.20E-07 | 1.70E-06 | 1.58 | 0.002 | 0.006 | 0.68 | | | 0.57 | | | 0.53 | | |
| **Ctsl** | 2.58 | 4.50E-12 | 1.50E-10 | 2.02 | 2.00E-09 | 1.40E-08 | 0.84 | 0.007 | 0.01 | 0.65 | | | 0.18 | | | -0.04 | | |
| **Junb** | 2.59 | 4.40E-08 | 2.60E-07 | 2.17 | 4.90E-07 | 1.40E-06 | 1.63 | 9.50E-05 | 0.0005 | 1.01 | 0.02 | 0.1 | 1.12 | 0.005 | 0.08 | 0.76 | | |
| **Rps6ka1** | 1.87 | 1.00E-05 | 3.60E-05 | 1.59 | 3.50E-05 | 7.40E-05 | 1.25 | 0.0009 | 0.002 | 0.91 | 0.02 | 0.1 | 1.23 | 0.001 | 0.03 | 0.31 | | |
| **Camkk1** | 0.18 | | | 0.78 | 0.002 | 0.003 | 0.7 | 0.005 | 0.01 | 0.52 | | | 0.39 | | | 0.41 | | |
| **Grb10** | -2.69 | 1.10E-16 | 1.50E-14 | -0.74 | 0.004 | 0.006 | -0.35 | | | -0.3 | | | -0.56 | 0.03 | 0.1 | -0.4 | | |
| **Akt2** | -1.4 | 3.30E-08 | 2.00E-07 | -1.38 | 2.70E-11 | 2.90E-10 | -0.77 | 0.0002 | 0.0009 | -0.45 | | | -0.49 | 0.02 | 0.1 | -0.26 | | |
| **Rps6kb1** | -1.59 | 5.50E-11 | 9.50E-10 | -1.47 | 8.80E-14 | 1.30E-12 | -0.7 | 0.0004 | 0.001 | -0.3 | | | -0.5 | 0.01 | 0.1 | -0.33 | | |
| **Eif4e** | -0.63 | 0.0001 | 0.0002 | -1.36 | 0 | 0 | -0.8 | 4.90E-09 | 9.60E-08 | -0.47 | 0.003 | 0.03 | -0.4 | 0.003 | 0.07 | -0.19 | | |
| **Eif2b5** | -0.93 | 0.0002 | 0.0005 | -1.26 | 2.90E-09 | 1.90E-08 | -1 | 2.10E-06 | 2.90E-05 | -0.49 | 0.04 | 0.1 | -0.35 | | | -0.2 | | |
| **Eif4g1** | -0.95 | 0.0004 | 0.0007 | -1.15 | 3.20E-07 | 9.80E-07 | -0.83 | 0.0002 | 0.0009 | -0.33 | | | -0.21 | | | -0.11 | | |
| **Rheb** | -0.81 | 6.60E-05 | 0.0001 | -1.12 | 3.40E-11 | 3.40E-10 | -0.75 | 1.10E-05 | 0.0001 | -0.44 | 0.02 | 0.1 | -0.38 | | | -0.21 | | |
| **Eif4b** | -0.96 | 0.0001 | 0.0003 | -1.21 | 1.10E-08 | 5.70E-08 | -0.73 | 0.0006 | 0.001 | -0.37 | | | -0.05 | | | 0.07 | | |
| **Mtor** | -1.1 | 4.30E-05 | 0.0001 | -0.83 | 0.0002 | 0.0003 | -0.63 | 0.004 | 0.008 | -0.24 | | | -0.07 | | | 0.02 | | |
| **Eif4ebp1** | 0.49 | 0.01 | 0.02 | -0.25 | | | -0.66 | 0.0001 | 0.0007 | -0.28 | | | -0.07 | | | 0.02 | | |
| **Akt1** | -0.29 | | | -0.39 | | | -0.44 | 0.0002 | 0.0009 | -0.02 | | | -0.37 | | | -0.2 | | |
| **Foxo1** | -1.4 | 1.50E-06 | 5.90E-06 | -1.22 | 3.70E-07 | 1.00E-06 | -0.6 | 0.01 | 0.01 | -0.15 | | | -0.19 | | | -0.18 | | |
| **Trim63** | -0.39 | | | -1 | 0.01 | 0.01 | -1.11 | 0.005 | 0.01 | -0.5 | | | -0.24 | | | -0.05 | | |
| **Fbxo32** | -0.37 | | | -0.13 | | | -0.37 | | | -0.33 | | | 0.72 | | | 0.32 | | |
| **Foxo3** | -0.85 | 0.0001 | 0.0003 | -0.07 | | | -0.15 | | | 0.3 | | | 0.15 | | | 0.01 | | |
| **Fst** | -0.91 | 0.005 | 0.008 | -0.32 | | | -0.65 | 0.01 | 0.02 | -0.03 | | | -0.4 | | | -0.38 | | |
| **Mstn** | -3.08 | 0.0001 | 0.0003 | -2.97 | 2.20E-06 | 5.60E-06 | -2.32 | 0.0001 | 0.0008 | -1.69 | 0.02 | 0.1 | -1 | | | -0.5 | | |
| **Ikbkb** | -0.95 | 9.40E-05 | 0.0002 | -1.14 | 1.50E-08 | 7.20E-08 | -0.78 | 0.0001 | 0.0005 | -0.53 | 0.02 | 0.1 | 0.41 | | | 0.33 | | |
| **Gsk3b** | -0.96 | 8.30E-05 | 0.0002 | -1.06 | 2.00E-07 | 6.60E-07 | -0.64 | 0.001 | 0.003 | -0.06 | | | 0.44 | | | -0.19 | | |
| **Nfkbia** | 0.47 | | | -0.27 | | | 0.23 | | | -0.02 | | | 0.57 | | | 0.53 | | |
| **Nol3** | -0.48 | | | -0.86 | 0.002 | 0.004 | -0.54 | | | -0.02 | | | 0.18 | | | -0.04 | | |
| **Prkaa1** | -0.29 | | | -0.34 | | | -0.27 | | | 0.18 | | | 1.12 | | | 0.76 | | |

**Table S5. Extracellular matrix remodeling related genes throughout the different time points after simulated birth injury.** P and q values are indicated for significantly differentially expressed genes compared to uninjured group. Log2FC

| | Days after Simulated Birth Injury | | | | | | | | | | | | | | | | | |
|---|---|---|---|---|---|---|---|---|---|---|---|---|---|---|---|---|---|---|
| **Gene** | 1 | | | 3 | | | 7 | | | 10 | | | 31 | | | 35 | | |
| | FC | p | q | FC | p | q | FC | p | q | FC | p | q | FC | p | q | FC | p | q |
| **Mmp8** | 2.74 | 0.002 | 0.004 | 0.84 | | | 1.58 | 0.04 | 0.06 | 0.47 | | | 0.33 | | | 0.6 | | |
| **Mmp9** | 1.32 | 0.02 | 0.03 | 1.79 | 0.001 | 0.001 | 1.66 | 0.002 | 0.005 | 1.38 | 0.01 | 0.1 | 0.28 | | | -0.41 | | |
| **Mmp14** | -0.87 | | | 1.29 | 0.001 | 0.003 | 1.75 | 5.50E-05 | 0.0003 | 1.05 | 0.02 | 0.1 | 0.51 | | | -0.01 | | |
| **Mmp2** | -2.35 | 1.60E-09 | 1.70E-08 | 0.49 | | | 1.39 | 4.70E-05 | 0.0003 | 1.13 | 0.002 | 0.02 | 0.33 | | | 0.14 | | |
| **Mmp1** | -1.6 | 0.0009 | 0.001 | -0.85 | | | -0.39 | | | -1.46 | 0.008 | 0.053 | -0.52 | | | -0.06 | | |
| **Timp1** | 4.28 | 7.20E-09 | 5.50E-08 | 4.46 | 4.70E-10 | 4.10E-09 | 2.66 | 3.60E-05 | 0.0002 | 1.28 | | | 1.25 | 0.03 | 0.1 | 0.44 | | |
| **Timp2** | -1.57 | 4.80E-11 | 9.50E-10 | -0.22 | | | 0.43 | 0.03 | 0.05 | 0.49 | 0.03 | 0.1 | 0.34 | | | 0.28 | | |
| **Timp3** | -2.26 | 2.60E-09 | 2.20E-08 | -1.55 | 2.60E-07 | 8.30E-07 | -0.15 | | | 0.84 | 0.02 | 0.1 | 0.54 | | | 0.61 | | |
| **Timp4** | -3.56 | 3.10E-12 | 1.40E-10 | -2.03 | 1.10E-07 | 4.20E-07 | -1.02 | 0.008 | 0.01 | -0.26 | | | 0.08 | | | -0.004 | | |
| **Col1a1** | -3.03 | 1.30E-05 | 4.30E-05 | 1.75 | 0.003 | 0.005 | 1.66 | 0.005 | 0.01 | 1.4 | 0.03 | 0.1 | -0.35 | | | -1.15 | 0.03 | 0.4 |
| **Col3a1** | -1.82 | 0.0009 | 0.001 | 1.92 | 0.0001 | 0.0002 | 2.08 | 5.10E-05 | 0.0003 | 1.66 | 0.002 | 0.02 | 0.36 | | | -0.19 | | |
| **Itgb1** | 0.39 | 0.04 | 0.05 | 0.35 | 0.04 | 0.05 | 0.18 | | | 0.42 | 0.03 | 0.1 | 0.02 | | | 0.007 | | |
| **Itga2** | -0.35 | | | 0.16 | | | -0.1 | | | -0.05 | | | 0.075 | | | -0.517 | | |
| **Itga1** | -1.76 | 4.40E-11 | 9.50E-10 | -0.16 | | | 0.24 | | | 0.52 | 0.04 | 0.1 | 0.21 | | | 0.02 | | |
| **Tgfbr1** | -0.27 | | | 0.28 | 0.0002 | 0.0003 | -0.01 | | | -0.35 | | | -0.06 | | | -0.07 | | |
| **Tgfb1** | 1.84 | 3.50E-05 | 0.0001 | 2.15 | 2.60E-07 | 8.30E-07 | 1.74 | 1.80E-05 | 0.0001 | 1.44 | 0.001 | 0.01 | 0.85 | 0.02 | 0.1 | 0.36 | | |
| **Smad3** | -1.19 | 0.006 | 0.009 | -2.02 | 1.50E-08 | 7.20E-08 | -1.44 | 5.30E-05 | 0.0003 | -1.18 | 0.006 | 0.04 | -0.71 | 0.04 | 0.1 | -0.49 | | |
| **Smad2** | -2.86 | | | -0.44 | | | -0.3 | | | 0.18 | | | 0.03 | | | -0.55 | | |
| **Smad4** | -0.92 | 2.80E-07 | 1.50E-06 | -0.89 | 3.70E-09 | 2.30E-08 | -0.5 | 0.0008 | 0.002 | -0.29 | | | -0.06 | | | 0.01 | | |
| **Pdfgra** | -0.06 | | | 1.06 | 4.00E-05 | 8.40E-05 | 0.5 | 0.04 | 0.06 | 0.45 | | | 0.41 | | | 0.44 | | |
| **Pdgfa** | -0.8 | 1.30E-05 | 4.30E-05 | -0.99 | 1.00E-10 | 9.40E-10 | -0.56 | 0.0002 | 0.0009 | -0.35 | | | -0.42 | 0.007 | 0.08 | -0.21 | | |
| **Vwa1** | 0.48 | | | 1.39 | 6.10E-06 | 1.40E-05 | 0.92 | 0.002 | 0.005 | 0.72 | 0.03 | 0.1 | 0.74 | 0.01 | 0.1 | 0.14 | | |

**Table S6. Vascularization related genes throughout the different time points after simulated birth injury.** P and q values are indicated for significantly differentially expressed genes compared to uninjured group. Log2FC

| | Days after Simulated Birth Injury | | | | | | | | | | | | | | | | | |
|---|---|---|---|---|---|---|---|---|---|---|---|---|---|---|---|---|---|---|
| **Gene** | 1 | | | 3 | | | 7 | | | 10 | | | 31 | | | 35 | | |
| | FC | p | q | FC | p | q | FC | p | q | FC | p | q | FC | p | q | FC | p | q |
| **Thsb1** | 4.7 | 4.00E-10 | 5.00E-09 | 3.6 | 9.80E-08 | 3.60E-07 | 2 | 0.0007 | 0.002 | 1.46 | 0.02 | 0.1 | 0.87 | | | 0.25 | | |
| **Fgf2** | -2.68 | 5.30E-07 | | -3.2 | 4.40E-14 | 7.80E-13 | -1.3 | 0.0009 | 0.002 | -1.35 | 0.007 | 0.05 | -0.83 | 0.04 | 0.18 | -0.48 | | |
| **Vegfa** | -2 | 2.00E-09 | 2.00E-08 | -2.54 | 0 | 0 | -1.78 | 1.10E-10 | 3.20E-09 | -1.27 | 0.0001 | 0.002 | -1.09 | 8.10E-05 | 0.003 | -0.71 | 0.01 | 0.25 |
| **Mmp14** | -0.87 | | | 1.29 | 0.002 | 0.003 | 1.75 | 5.50E-05 | 0.0003 | 1.05 | 0.02 | 0.1 | 0.51 | | | -0.01 | | |
| **Nrp1** | -0.14 | | | 0.3 | | | 0.26 | | | 0.41 | | | 0.1 | | | -0.25 | | |
| **Tie1** | -0.3 | | | 0.76 | 0.0001 | 0.0002 | 0.64 | 0.001 | 0.003 | 0.68 | 0.002 | 0.02 | 0.42 | 0.03 | 0.14 | 0.2 | | |
| **Angpt2** | -0.95 | 0.0001 | 0.0004 | -0.42 | 0.04 | 0.06 | 0.05 | | | -0.04 | | | 0.33 | | | 0.28 | | |
| **Flt1** | -0.69 | 0.009 | 0.01 | -0.74 | 0.001 | 0.001 | -0.57 | 0.01 | 0.01 | -0.2 | | | -0.05 | | | -0.22 | | |
| **Kdr** | -1.05 | 0.0002 | 0.0004 | -0.47 | 0.04 | 0.06 | -0.14 | | | -0.1 | | | -0.31 | | | -0.83 | 0.0006 | 0.02 |
| **Angpt1** | -2.12 | 1.00E-13 | 7.20E-12 | -1.35 | 1.50E-09 | 1.10E-08 | -0.45 | 0.04 | 0.06 | -0.006 | | | -0.29 | | | 0.001 | | |
| **Vegfb** | -1.62 | 1.10E-06 | 4.60E-06 | -2 | 3.60E-14 | 7.80E-13 | -0.98 | 0.0003 | 0.001 | -0.39 | | | -0.59 | 0.03 | 0.14 | -0.46 | | |

**Table S7. Neuromuscular junctions related genes throughout the different time points after simulated birth injury.** P and q values are indicated for significantly differentially expressed genes compared to uninjured group. Log2FC

| | Days after Simulated Birth Injury | | | | | | | | | | | | | | | | | |
|---|---|---|---|---|---|---|---|---|---|---|---|---|---|---|---|---|---|---|
| **Gene** | 1 | | | 3 | | | 7 | | | 10 | | | 31 | | | 35 | | |
| | FC | p | q | FC | p | q | FC | p | q | FC | p | q | FC | p | q | FC | p | q |
| **Dvl1** | -1.58 | 2.10E-10 | 3.00E-09 | -1.69 | 1.10E-16 | 3.80E-15 | -0.77 | 0.0001 | 0.0007 | -0.3 | | | -0.43 | 0.03 | 0.14 | -0.16 | | |
| **Gphn** | -1.34 | 1.70E-06 | 6.60E-06 | -1.4 | 1.20E-09 | 9.80E-09 | -0.79 | 0.0006 | 0.001 | -0.38 | | | -0.23 | 0.32 | 0.55 | -0.05 | | |
| **Utrn** | -1.79 | 3.50E-09 | 2.90E-08 | -0.63 | 0.01 | 0.01 | -0.12 | | | 0.21 | | | 0.15 | | | -0.04 | | |
| **Agrn** | -1.16 | 0.0004 | 0.0007 | 0.04 | | | 0.43 | | | 0.47 | | | 0.25 | | | -0.19 | | |
| **App** | -0.78 | 0.0002 | 0.0004 | 0.29 | | | 0.43 | 0.01 | 0.02 | 0.62 | 0.003 | 0.03 | 0.22 | | | -0.1 | | |
| **Tnc** | 1.61 | 0.03 | 0.04 | 3.93 | 6.90E-07 | 1.90E-06 | 2.82 | 0.0001 | 0.0007 | 1.69 | 0.02 | 0.1 | -0.1 | | | -0.85 | | |
| **Nrg1** | 1.58 | 0.008 | 0.01 | 1.33 | 0.01 | 0.01 | 1.12 | 0.03 | 0.05 | -0.002 | | | 0.03 | | | 0.12 | | |
| **Ptn** | -1.18 | 0.01 | 0.02 | 2.08 | 5.80E-06 | 1.30E-05 | 1.91 | 2.50E-05 | 0.0002 | 0.96 | 0.04 | 0.12 | 0.58 | | | 0.38 | | |
| **Erbb2** | 0.06 | | | 1.43 | 0.0001 | 0.0002 | 1.24 | 0.001 | 0.002 | 1.72 | 3.90E-05 | 0.0009 | 0.95 | 0.01 | 0.1 | 0.23 | | |
| **Musk** | -0.52 | | | 1.84 | 4.60E-06 | 1.00E-05 | 1.48 | 0.0002 | 0.0008 | 2.05 | 3.20E-06 | 8.80E-05 | -0.21 | | | -0.24 | | |

**Table S8. Composition of decellularized porcine skeletal muscle ECM.**

| *Protein Abundance in Porcine Skeletal Muscle Matrix (nmol/g)* | | | |
|---|---|---|---|
| **Protein** | **Gene** | **Functional Classification** | **Average** |
| Collagen alpha-1(IV) chain(Arresten/Core Protein) | COL4A1 | Basement Membrane | 25.55 |
| Collagen alpha-1/5(IV) chain(Arresten/Core Protein) | COL4A1/5 | Basement Membrane | 92.89 |
| Collagen alpha-2(IV) chain(Canstatin/Core Protein) | COL4A2 | Basement Membrane | 19.35 |
| Laminin Beta-1 | LAMB1 | Basement Membrane | 0.15 |
| Laminin Beta-2 | LAMB2 | Basement Membrane | 0.17 |
| Laminin Gamma-1 | LAMC1 | Basement Membrane | 0.21 |
| Perlecan | HSPG2 | Basement Membrane | 2.99 |
| Collagen alpha-1(XIV) chain | COL14A1 | FACIT Collagen | 0.40 |
| Collagen alpha-1(I) chain | COL1A1 | Fibrillar Collagen | 1393.12 |
| Collagen alpha-1(V) chain | COL5A1 | Fibrillar Collagen | 35.55 |
| Collagen alpha-2(I) chain | COL1A2 | Fibrillar Collagen | 1272.17 |
| Collagen alpha-2(V) chain | COL5A2 | Fibrillar Collagen | 39.68 |
| Collagen alpha-1(VI) chain | COL6A1 | Matricellular | 28.34 |

| | | | |
|---|---|---|---|
| Collagen alpha-2(VI) chain | COL6A2 | Matricellular | 3.36 |
| Collagen alpha-3(VI) chain | COL6A3 | Matricellular | 23.05 |
| Dermatopontin | DPT | Matricellular | 1.30 |
| Emilin 1 | EMILIN1 | Matricellular | 0.22 |
| Fibronectin 1(type-III 4 domain) | FN1 | Matricellular | 4.53 |
| Lumican | LUM | Matricellular | 13.65 |
| Fibrillin 1 | FBN1 | Structural ECM | 3.12 |
| Actin (All Isoforms) | ACT | Cytoskeletal | 9.00 |
| Myosin(Myosin-3,4,6,7) | MYH | Cytoskeletal | 3.12 |

Note, low cytoskeletal components indicate sufficient decellularization.
